# A genome-wide overexpression screen identifies *Tb*FOP, whose overexpression disrupts transcription termination in *Trypanosoma brucei*

**DOI:** 10.1101/2025.09.02.673782

**Authors:** Navina Panneer Selvan, Aditi Mukherjee, Vivek Gupta, Esteban Erben, Hee-Sook Kim

## Abstract

*Trypanosoma brucei* is a protozoan parasite that causes African trypanosomiasis. Understanding essential cellular processes in *T. brucei* can provide insights into parasite-specific vulnerabilities. We therefore screened for genes whose overexpression impaired parasite growth. Three top-ranking candidates were confirmed to inhibit *T. brucei* growth upon overexpression. Among them, *Tb*FOP, a previously uncharacterized protein, exhibited distinct phenotypes beyond growth inhibition. *Tb*FOP overexpression caused severe cell-cycle defects, substantially increasing the proportion of sub-G1 cells lacking nuclear DNA. Transcriptome analysis revealed widespread transcriptional dysregulation following *Tb*FOP overexpression. Specifically, antisense transcript levels substantially increased near transcription termination sites (TTSs) across the genome. *Tb*FOP overexpression also increased the expression of normally silent variant surface glycoprotein (VSG) genes across multiple genomic contexts. These phenotypes resemble those observed in trypanosome mutants lacking TTS-associated chromatin marks, including two histone variants and base J, a kinetoplastid-specific DNA modification. The preferential effects at TTSs and VSG loci raise the possibility that *Tb*FOP overexpression interferes with chromatin-associated transcriptional regulation at specific genomic regions. These findings demonstrate that *Tb*FOP overexpression profoundly affects transcription, cell-cycle progression, and parasite growth.

**IMPORTANCE:** *Trypanosoma brucei*, the parasite that causes African trypanosomiasis, survives in mammalian hosts by periodically changing its major surface protein to evade immune responses. While this process is essential for parasite survival and requires strict control of gene expression, many of the underlying regulatory mechanisms remain poorly understood. Here, a genome-wide toxicity screen identified *Tb*FOP along with other proteins whose overproduction impairs parasite growth. *Tb*FOP is a previously uncharacterized protein whose overproduction severely disrupts transcriptional regulation as well as parasite growth. Increased *Tb*FOP levels caused widespread transcriptional dysregulation, including increased antisense transcript levels near transcription termination sites and inappropriate expression of normally silent surface protein genes. These findings reveal a previously unrecognized link among altered *Tb*FOP expression, transcriptional regulation, and VSG gene silencing. Understanding these unusual regulatory processes is important for defining how trypanosomes maintain persistent infections and may reveal biological processes that distinguish these parasites from their mammalian hosts.

## INTRODUCTION

*Trypanosoma brucei* is a protozoan parasite that causes African trypanosomiasis in humans (human African trypanosomiasis, HAT) and animals (animal African trypanosomiasis, AAT) throughout sub-Saharan Africa. Trypanosome parasites are transmitted to mammalian hosts through the bite of an infected tsetse fly. In the mammalian bloodstream, the parasite differentiates into the bloodstream form (BF), which is characterized by a dense surface coat composed of a single variant surface glycoprotein (VSG) species. A single VSG gene is expressed from a repertoire of approximately 2,500 VSG genes and pseudogenes dispersed throughout the parasite genome. The VSG coat triggers a strong adaptive immune response in the host and plays an essential role in the parasite’s immune evasion. *T. brucei* periodically changes the monoallelically selected VSG allele through transcriptional and recombination mechanisms, a process known as VSG switching [1–4]. Through monoallelic VSG expression and VSG switching, the parasite persists in the host, causing death if left untreated.

The *T. brucei* genome harbors 11 pairs of megabase-sized chromosomes, which are organized into three regions: the RNA polymerase II (Pol II)-transcribed core, transcriptionally inactive subtelomeric regions, and the RNA Pol I-transcribed telomeric VSG expression site (ES), which houses a VSG gene [5, 6]. Unlike those of most eukaryotes, *T. brucei* genes are assembled into polycistronic transcription units (PTUs)[7–10]. PTUs are delimited by transcription start sites (TSSs) and transcription termination sites (TTSs), which are marked by specific histone modifications and histone variants [11, 12]. Histone H4K10ac, H3K4me^3^, and two essential histone variants (H2Az and H2Bv) are enriched at TSSs, whereas two non-essential histone variants (H3v and H4v) are present at TTSs. In addition, TSS and TTS DNA contain a kinetoplastid-specific DNA modification, base J (β-D-glucosyl-5-hydroxymethyluracil) [13]. Base J is synthesized in two steps: first, hydroxylation of dT (5mU) to 5-hydroxymethyl-dU (5hmU) by J-binding proteins 1 and 2 (JBP1 and JBP2), and then glucosylation of 5hmU to glucosyl-5hmU (base J) by J-glucosyl transferase (JGT) [14, 15].

Base J plays a crucial role in transcription termination in trypanosomatids, including *T. brucei* and *Leishmania*, either alone or in cooperation with histone variants [16–19]. However, the molecular mechanism linking base J to transcription termination remains incompletely understood. The identification of J-binding protein 3 (JBP3) provided an important mechanistic connection between base J and the transcription termination machinery. In *Leishmania* and *T. brucei*, JBP3 was identified through the affinity purification of JGT, which led to the identification of the transcription termination complex PJW/PP1, consisting of PNUTS (PP1 nuclear targeting subunit), JBP3, WDR82 (WD40 repeat protein), and PP1-1 (protein phosphatase 1-1) [20–23]. Depletion of components of PJW/PP1 resulted in cell lethality and readthrough transcription at TTSs in *Leishmania* and *T. brucei* [20–23].

Base J is enriched at two other loci in the *T. brucei* genome, telomere repeats and VSG ESs [24, 25]. Only one VSG gene is transcribed among thousands of VSG alleles distributed in four different genomic environments. Two loci, the bloodstream expression site (BES) [26, 27] and the metacyclic expression site (MES) [28, 29], each harbor an RNA Pol I promoter and a VSG gene upstream of the telomere repeats. BESs contain additional genes, such as expression-site-associated genes (ESAGs), while MESs have only a VSG gene. About 15 BESs and 5–6 MESs are located upstream of megabase chromosome telomere repeats. Additional VSG genes reside on about 50 minichromosomes (MC VSGs) [29, 30] or are dispersed in arrays in subtelomeric regions (chromosome-internal, CI VSGs) [6]. MC and CI VSGs lack promoters and are naturally silenced [30]. Base J is enriched in BESs, especially at repeat sequences, including the 50-bp repeat situated upstream of the promoter and the 70-bp repeats located upstream of the VSG gene. Depletion of *T. brucei* PP1-1 of the PJW/PP1 complex or depletion of TTS chromatin marks disrupted the BES silencing, increasing the levels of silent VSG and ESAG transcripts [23]. Therefore, base J biosynthesis and PJW/PP1 components are involved not only in regulating RNA Pol II transcription termination but also in controlling VSG expression by Pol I.

Many essential cellular processes in *T. brucei*, including transcription termination, have diverged from those in mammalian hosts, providing opportunities to identify parasite-specific vulnerabilities. To gain insight into essential cellular processes in *T. brucei*, we performed a genome-wide overexpression-induced toxicity (OE toxicity) screen using the ORFeome libraries [31]. Screening approximately 400,000 transfectants identified about 200 candidate genes, from which three top-ranking candidates, *Tb*927.6.1470 (*Tb*FOP), *Tb*927.5.1270, and *Tb*927.11.2250, were validated for growth inhibition upon overexpression. Further characterization of *Tb*FOP revealed that its overexpression increases antisense transcription at TTSs and disrupts VSG gene repression. Here, we report the results of the screen and characterize the phenotypes associated with *Tb*FOP overexpression.

## RESULTS

### Genome-wide overexpression-induced toxicity screen in BF *T. brucei*

To identify factors involved in essential cellular processes in *T. brucei*, we performed a genome-wide overexpression-induced toxicity screen using the *T. brucei* ORFeome libraries [31]. We used a WT BF *T. brucei* strain in the 2T1 background (AMT2), in which a specific rDNA locus contains a landing pad (LP) consisting of a puromycin-resistance gene (PUR) and the C-terminal portion of a hygromycin-resistance gene (HYG) [31, 32]. The AscI-linearized pRPa-Sce* vector containing a Tet-inducible I-SceI endonuclease gene, its 18-bp recognition site, and an overlapping N-terminal portion of HYG was integrated into the LP by homologous recombination (Fig. S1). Correct integration replaces PUR and reconstitutes a functional HYG gene; therefore, hygromycin-resistant and puromycin-sensitive clones were selected for subsequent transfection of the ORFeome library. Tetracycline induction of I-SceI generates a DSB at the LP, facilitating integration of the ORFeome library at the same locus. After 3 hours of Tet induction, cells were transfected with the ORFeome library vectors linearized by I-SceI [31]. Approximately 400,000 clones were obtained.

Triplicate cultures with or without tetracycline were grown for 3 days, and the representation of each ORF in Tet-treated cultures relative to untreated cultures was analyzed by targeted high-throughput sequencing. Genomic DNA isolated from each culture was fragmented, ligated to an Illumina adaptor, and PCR-amplified using a custom p5 forward primer that anneals to the attB1 site of the ectopic ORF and an indexed p7 primer, as described previously [31]. The resulting PCR amplicons were analyzed by high-throughput sequencing on an Illumina platform.

To identify ORFs whose overexpression inhibits *T. brucei* growth (underrepresented in Tet-treated cultures), sequence reads were aligned to the *Tb*927v5 genome using Bowtie 2 and analyzed with SeqMonk and DESeq (Fig. 1A and Table S5). We identified 193 underrepresented ORFs in Tet-treated cultures (*P* adj < 0.1), representing candidate genes whose overexpression impairs *T. brucei* growth (Table S5D). These 193 underrepresented ORFs were categorized according to their known or predicted functions (Fig. 1B). Candidates were diverse: 64% (124 of 193 ORFs) had known or predicted functions, whereas 36% (69) were annotated as hypothetical genes (HYP genes). Among the genes with known or predicted functions, approximately 29% (36 of 124 ORFs) appear to be involved in RNA metabolism, including RNA binding, degradation, and processing. Hypothetical genes were further classified using the subcellular localization dataset [33]. Of these, 17% (12 of 69 ORFs) showed nuclear localization, whereas no localization data were available for about 55% of the HYP genes (38 of 69 ORFs).

**FIG 1.**
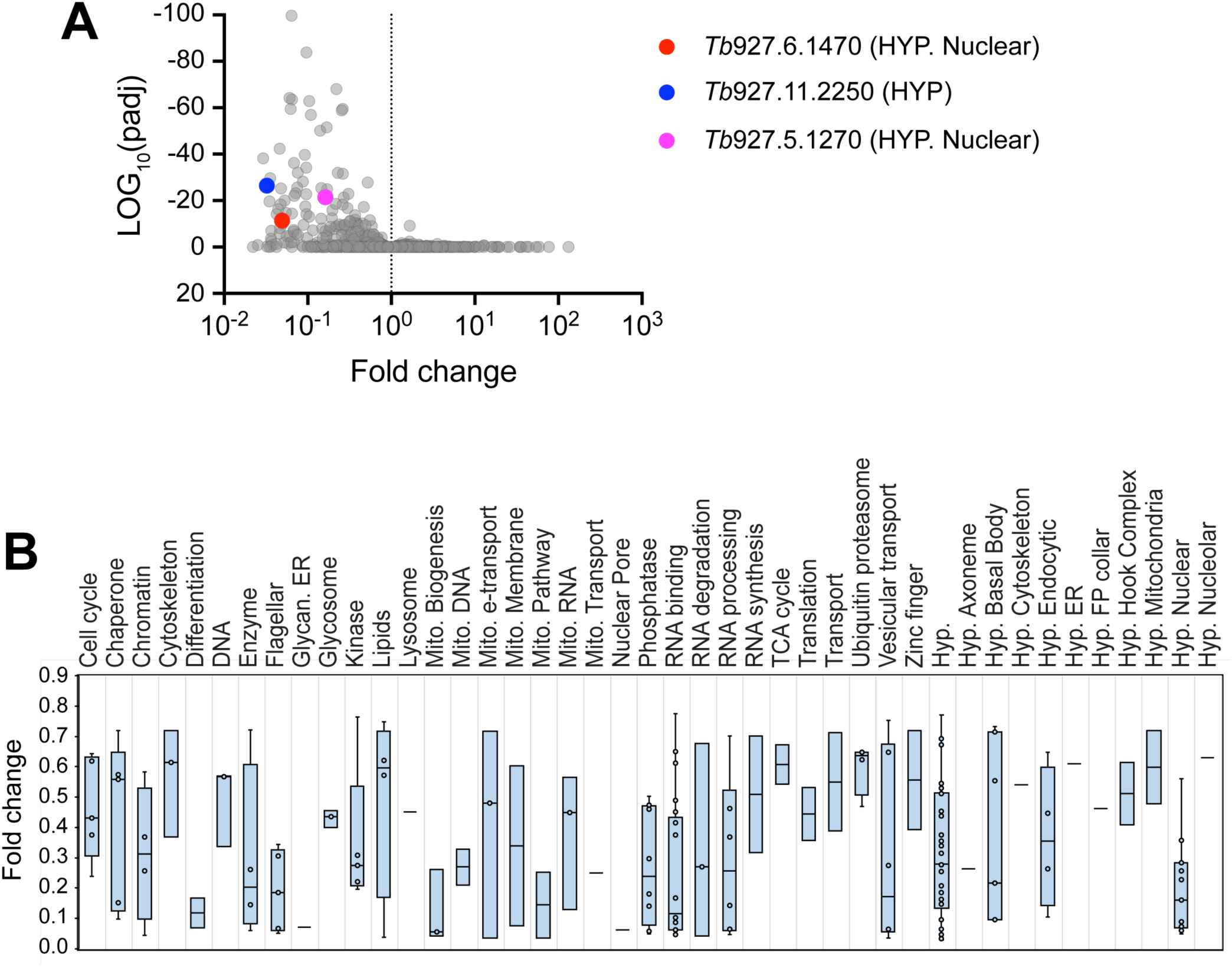
Identification of genes whose overexpression inhibits *T. brucei* cell growth. **(A)** Overexpression-induced toxicity screen. Wild-type (WT) bloodstream-form *T. brucei* cells transfected with the ORF library were screened for overexpression-induced growth inhibition. ORF-transfected cells were cultured with or without tetracycline (Tet) in three independent cultures for 3 days. ORFs whose overexpression caused toxicity were expected to be underrepresented in Tet-treated cultures. Candidate toxic ORFs were identified using targeted next-generation sequencing (NGS) and DESeq analysis (see Fig. S1 and Table S5 for details). Three candidates, *Tb*927.6.1470 (*Tb*FOP, red), *Tb*927.11.2250 (blue), and *Tb*927.5.1270 (pink), were selected for further validation. **(B)** Candidate classification. Candidates were categorized according to their predicted functions and subcellular localizations reported in TrypTag [33] (http://tryptag.org/). Underrepresented candidates with adjusted *P* values (*P* adj) < 0.1 are shown; each circle represents one ORF. See Table S5 for details.

Consistent with the ability of the screen to recover growth-inhibitory ORFs, several genes previously reported to induce toxicity were identified, including RNA-binding protein 10 (RBP10, fold change (FC) = 0.06, *P* adj = 3 x 10^-60^) and RBP9 (FC = 0.12, *P* adj = 6.5 x 10^-8^). Both are involved in *T. brucei* differentiation from the bloodstream form to the insect-stage procyclic form (PF) [34, 35]. The screen also identified VEX2 (FC = 0.32, *P* adj = 1.1 x 10^-11^), which is crucial for monoallelic VSG expression and DOT1B (FC = 0.04, *P* adj = 5.6 x 10^-17^), a histone H3 methyltransferase responsible for H3K76 trimethylation and involved in genome integrity and antigenic variation [36–38]. However, DOT1A, another H3 methyltransferase known to inhibit *T. brucei* cell growth upon overexpression [39], was not identified. DOT1A was among the ∼1,600 ORFs that were not recovered from the ∼7,300 ORFs initially amplified, corresponding to an overall ORF recovery of about 78% (summarized in Table S5 C and F).

### Validation of three uncharacterized candidates: growth inhibition and modulation of BES silencing

Three top-ranking candidates with potential links to VSG expression regulation or *T. brucei* life-cycle differentiation were selected for validation: *Tb*927.11.2250 (FC = 0.03, *P* adj = 3.8 x 10^-27^), *Tb*927.6.1470 (FC = 0.05, *P* adj = 3.8 x 10^-12^), and *Tb*927.5.1270 (FC = 0.16, *P* adj = 3.2 x 10^-22^) (Fig. 1A, Table S5D).

*Tb*927.6.1470 contains a FOP (friends of PRMT1) domain found in several proteins involved in mRNA splicing and export. The *Trypanosoma cruzi* FOP protein was identified as one of the interacting proteins of *Tc*SUB2 and *Tc*eIF4AIII, RNA helicases involved in mRNA processing and export [40]. However, depletion of *Tb*927.6.1470 in *T. brucei* did not affect global mRNA export [40] and thus, its precise role in RNA metabolism remains unclear. In addition, *Tb*927.6.1470 was identified in the proteome of *Tb*RAP1 (repressor activator protein 1), which plays a central role in telomere protection and the regulation of monoallelic VSG expression [41, 42]. Hereafter, *Tb*927.6.1470 is referred to as *Tb*FOP.

*Tb*927.5.1270 is a putative member of the acidic leucine-rich nuclear phosphoprotein 32 (ANP32) family. In other eukaryotes, ANP32 proteins act as histone chaperones; for example, ANP32e regulates the abundance of soluble and chromatin-bound H2A.Z [43, 44]. ANP32B chaperones the histone variant macroH2A, which is involved in heterochromatin organization and transcriptional regulation [45]. If *Tb*927.5.1270 is an ANP32 ortholog, its overexpression could potentially perturb chromatin-associated silencing of ES-associated VSG genes.

*Tb*927.11.2250 was identified in the mRNA-bound proteome [46, 47] and as a regulatory component associated with BF *T. brucei* differentiation from the proliferative slender form to the non-proliferative stumpy form [48, 49].

To validate these genes for OE-induced toxicity, we first generated a reporter cell line containing a green fluorescent protein (GFP) reporter gene integrated downstream of a silent BES promoter (Fig. 2A). Tet-inducible genes encoding N-terminally 6xHis-tagged *Tb*FOP, *Tb*927.11.2250, or *Tb*927.5.1270 were integrated at the rDNA spacer LP locus. WT and OE-inducible strains were cultured with or without tetracycline and examined daily for cell growth (Fig. 2B and 2D) and viability (Fig. 2C and 2E). Consistent with the toxicity screen results, overexpression of *Tb*FOP and *Tb*927.11.2250 caused severe growth defects, with approximately 100-and 70-fold reductions in cell number by day 2, respectively (Fig. 2B). The alamarBlue assay [50, 51] showed 21.2% and 28.5% viability, respectively, for *Tb*FOP OE and *Tb*927.11.2250 OE cells on day 1; on day 2, viability decreased to 1.4% and 3.4%, respectively (Fig. 2C). These results confirm the severe growth-inhibitory effects of their overexpression. Because *Tb*927.5.1270 (FC = 0.16) was less strongly underrepresented in the screen than *Tb*FOP (FC = 0.05) or *Tb*927.11.2250 (FC = 0.03), we anticipated a milder growth phenotype. Consistent with this expectation, *Tb*927.5.1270 OE caused a modest effect on trypanosome cell growth (Fig. 2D and 2E).

**FIG 2.**
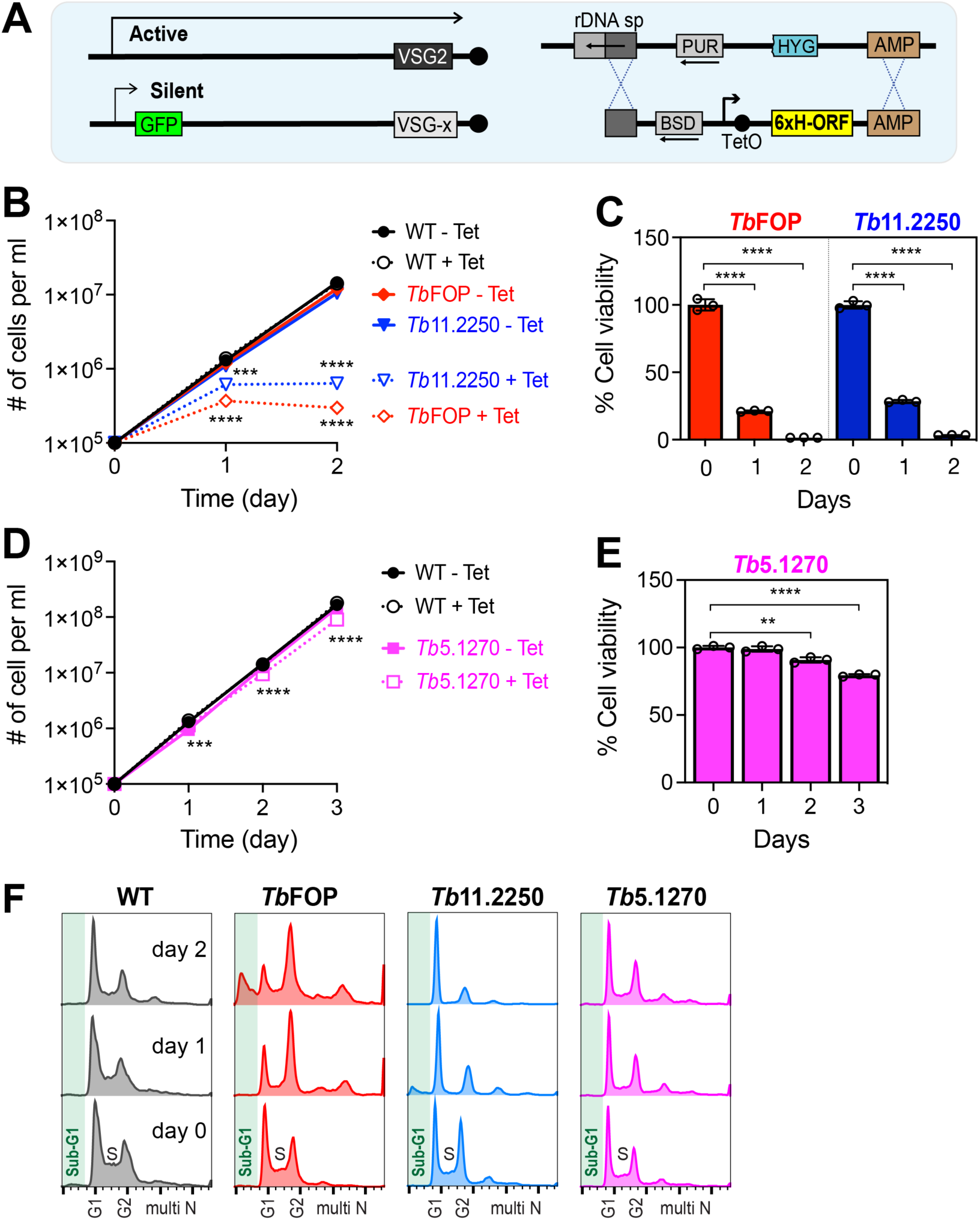
Validation of three candidates for overexpression-induced growth inhibition. **(A)** Validation strain containing a GFP reporter gene. Each ORF encoding N-terminally 6×His-tagged *Tb*FOP, *Tb*927.11.2250, or *Tb*927.5.1270 was individually integrated into the rDNA spacer region of a strain harboring a GFP reporter gene downstream of a silent BES promoter. Overexpression was induced by the addition of tetracycline. **(B, C)** Overexpression of *Tb*FOP or *Tb*927.11.2250 severely impairs *T. brucei* cell growth. *Tb*FOP and *Tb*927.11.2250 OE-inducible strains were cultured with or without Tet for 2 days. Cell growth and viability were monitored daily by cell counting (B) and AlamarBlue assays (C), respectively. **(D, E)** *Tb*927.5.1270 overexpression mildly inhibits *T. brucei* cell growth. The *Tb*927.5.1270 OE-inducible strain was analyzed as described for panels B and C for 3 days. Statistical significance was determined using two-tailed unpaired *t* tests: **, *P* < 0.01; ***, *P* < 0.001; and ****, *P* < 0.0001. Error bars represent the standard deviation (SD) of triplicate measurements (*n* = 3). Data are representative of three independent experiments. **(F)** Cell-cycle analysis. WT and OE-inducible strains were treated with Tet, fixed in ice-cold 75% ethanol, stained with propidium iodide, and analyzed by flow cytometry.

To examine whether overexpression of these genes also impairs cell cycle progression, WT and OE cells were stained with propidium iodide (PI) and cell-cycle profiles were analyzed by flow cytometry (Fig. 2F and Fig. S2). *Tb*FOP overexpression induced substantial cell-cycle defects, increasing G2/M, sub-G1, and other abnormal cell populations on days 1 and 2. Overexpression of *Tb*927.11.2250 caused a significant reduction in the S-phase population. As shown in Fig. S3, the percentage of cells in S phase decreased from 28.6% to 10.3% after Tet induction, whereas the percentage of G1 cells increased from 44.5% to 66.1%. The G2/M population remained unchanged. *Tb*927.5.1270 OE did not substantially alter cell-cycle progression (Fig. 2F).

Given the potential links of *Tb*FOP and *Tb*927.5.1270 to chromatin-associated processes [42–45], we examined whether overexpression of *Tb*FOP, *Tb*927.5.1270, or *Tb*927.11.2250 affects expression of the GFP reporter integrated downstream of a silent BES promoter. The expression of 6xHis-tagged *Tb*FOP, *Tb*927.11.2250, and *Tb*927.5.1270 was confirmed by immunoblotting (Fig. 3A). The level of VSG3, a normally silent VSG in our strain background in which VSG2 is actively expressed, increased in *Tb*FOP OE and *Tb*927.5.1270 OE cells. GFP levels also increased in these cells, indicating that overexpression of *Tb*FOP or *Tb*927.5.1270 perturbs BES silencing.

**FIG 3.**
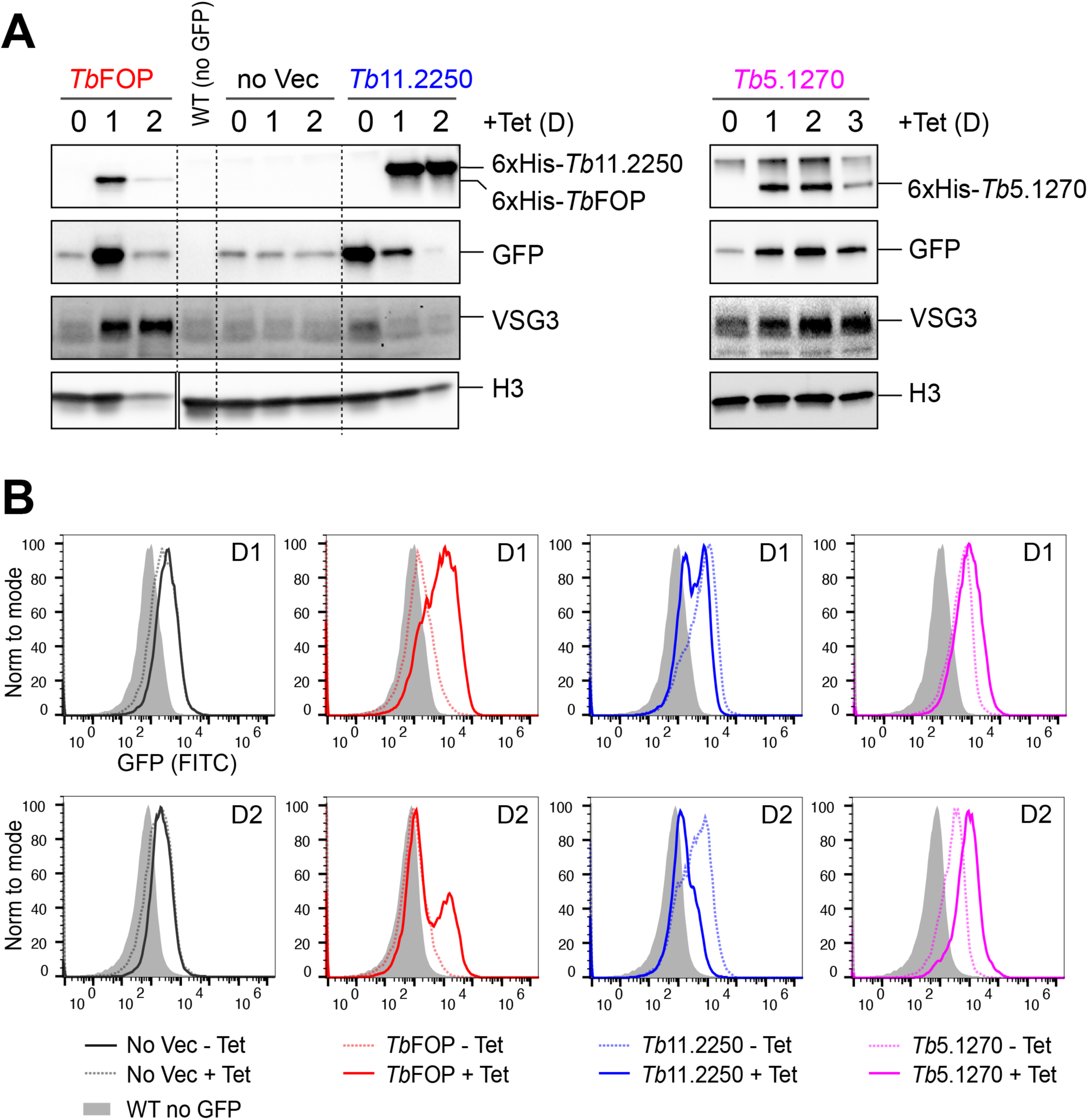
Overexpression of *Tb*FOP, *Tb*927.11.2250, and *Tb*927.5.1270 differentially modulates BES silencing. **(A)** Effects of candidate overexpression on silent BES-associated VSG3 and GFP protein levels. Overexpression of *Tb*FOP or *Tb*927.5.1270 increased the levels of VSG3 and GFP proteins, whereas overexpression of *Tb*927.11.2250 decreased their levels. Whole-cell lysates collected at the indicated time points were analyzed by immunoblotting using antibodies against the 6xHis, GFP, VSG3, or histone H3. Lysates corresponding to 2.5 × 10⁶ cells were loaded per lane. Histone H3 served as a loading control. **(B)** Flow-cytometric analysis of GFP expression. Live cells collected at the indicated time points were analyzed by flow cytometry. D1 and D2 indicate days 1 and 2 after Tet addition, respectively. No Vec denotes the parental GFP reporter strain lacking an integrated OE construct.

In contrast, *Tb*927.11.2250 OE decreased GFP and VSG3 protein levels. Together with the reduction in the S-phase population, the decreased VSG3 level, its previously reported involvement in slender-to-stumpy differentiation, and its identification in RNA-bound proteomes [46–49], these observations suggest that the phenotypes caused by *Tb*927.11.2250 overexpression may be related to pathways involved in stumpy formation or life-cycle differentiation.

Flow cytometry further confirmed the changes in GFP expression (Fig. 3B). Consistent with the immunoblotting data, GFP protein levels increased in Tet-treated *Tb*FOP OE cells on day 1 but decreased on day 2. This decrease is likely associated with the severe loss of cell viability and cell-cycle defects caused by *Tb*FOP OE on day 2, which were accompanied by reduced H3 levels. GFP levels increased over time following induction of *Tb*927.5.1270 OE. In contrast, *Tb*927.11.2250 OE decreased GFP levels, consistent with the immunoblotting data. Together, these results show that overexpression of the three candidate genes differentially affects silent BES expression.

### *Tb*FOP overexpression disrupts VSG silencing

Based on the pronounced effects of *Tb*FOP OE on BES silencing, we further characterized *Tb*FOP. To examine the effects of *Tb*FOP overexpression and depletion on VSG expression and global transcription, we performed rRNA-depleted stranded RNA-seq in *Tb*FOP-overexpressing and *Tb*FOP-depleted cells. One endogenous *Tb*FOP allele was tagged with 3xHA at the C-terminus in the GFP reporter strain, and a Tet-inducible *Tb*FOP knockdown (KD) construct was introduced (Fig. 2A). *Tb*FOP depletion did not significantly affect cell growth or BES silencing: GFP and VSG3 protein levels were unchanged upon *Tb*FOP depletion (Fig. S4).

To examine the effects of *Tb*FOP OE or KD on VSG expression, RNA-seq reads were aligned with Bowtie2 to VSGnome [30], a database containing sequences of ∼2,500 VSG genes distributed across four major chromosomal locations [6, 29, 30]. These include Pol I-transcribed BES and MES VSGs and promoterless MC and CI VSGs (Fig. 4A). The active VSG, VSG2, is located in BES1. Reads were analyzed using SeqMonk, and RPKM (reads per kilobase per million mapped reads) values were obtained for each VSG coding sequence. Log2-transformed RPKM values were compared between WT and *Tb*FOP OE cells (Fig. 4B) and between WT and *Tb*FOP KD cells (Fig. 4C). Approximately 27% of VSG genes were upregulated by at least 2-fold in *Tb*FOP-overexpressing cells (*q* < 0.05), whereas *Tb*FOP depletion did not significantly affect VSG expression.

**FIG 4.**
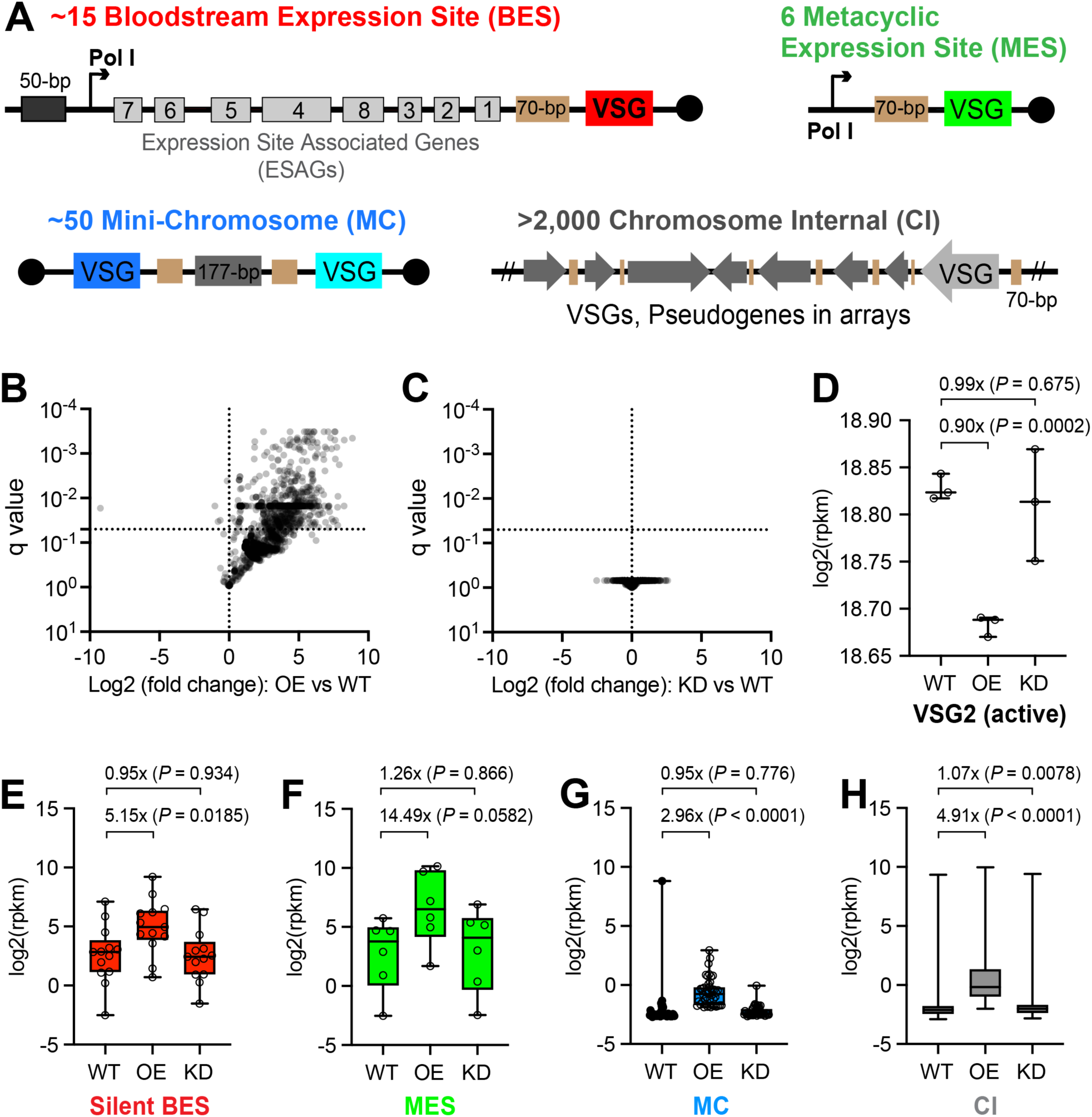
Overexpression of *Tb*FOP disrupts VSG silencing. **(A)** Diagrams illustrating four classes of genomic loci harboring VSG alleles: bloodstream expression sites (BESs), metacyclic expression sites (MESs), minichromosomes (MCs), and chromosome-internal (CI) loci. **(B, C)** Effects of *Tb*FOP overexpression and depletion on silent VSG transcript levels. rRNA-depleted stranded RNA-seq was performed using three independent cultures of each WT, *Tb*FOP-overexpressing, and *Tb*FOP-depleted strain. Sequence reads were aligned to the VSGnome database. Log2 fold changes and corresponding *q* values for individual VSG genes were plotted for OE versus WT (B) and KD versus WT (C). *P* values were calculated using multiple unpaired *t* tests with Welch’s correction, and *q* values were determined using the two-stage linear step-up procedure to control the false discovery rate in GraphPad Prism. The horizontal dotted line indicates *q* = 0.05. **(D–H)** Box plots showing the effects of *Tb*FOP OE and KD on different VSG classes. VSG2, the active VSG (D); silent BES VSGs (E); MES VSGs (F); MC VSGs (G); and CI VSGs (H) are shown. Statistical significance within each VSG class was determined using unpaired *t* tests with Welch’s correction in GraphPad Prism.

We next examined whether *Tb*FOP OE differentially affected specific classes of VSG genes by analyzing BES, MES, MC, and CI VSGs separately (Fig. 4D–H). *Tb*FOP OE increased the expression of ES-associated VSGs, with MES and BES VSG transcript levels increasing by ∼14.5-fold and ∼5.1-fold, respectively (Fig. 4E and 4F). Subtelomeric CI VSGs were also significantly upregulated, whereas MC VSGs showed the smallest increase. Together, these results show that *Tb*FOP overexpression perturbs the silencing of multiple classes of normally silent VSGs, with the strongest effects observed for ES-associated and CI VSGs. In contrast, *Tb*FOP depletion did not significantly alter the expression of either silent VSGs or the active VSG.

### *Tb*FOP overexpression increases antisense transcription across TTSs

To investigate whether *Tb*FOP OE or KD affects Pol II transcription, we separately analyzed forward and reverse RNA-seq reads mapping to the core regions of the 11 megabase chromosomes using 5-kb sliding windows with a 1-kb step (Fig. 5). Forward reads mapping to reverse PTUs (blue; transcribed from right to left) represent sense transcription, whereas forward reads mapping to forward PTUs (red; transcribed from left to right) represent antisense transcription. Conversely, reverse reads mapping to forward PTUs represent sense transcription, whereas reverse reads mapping to reverse PTUs represent antisense transcription.

**FIG 5.**
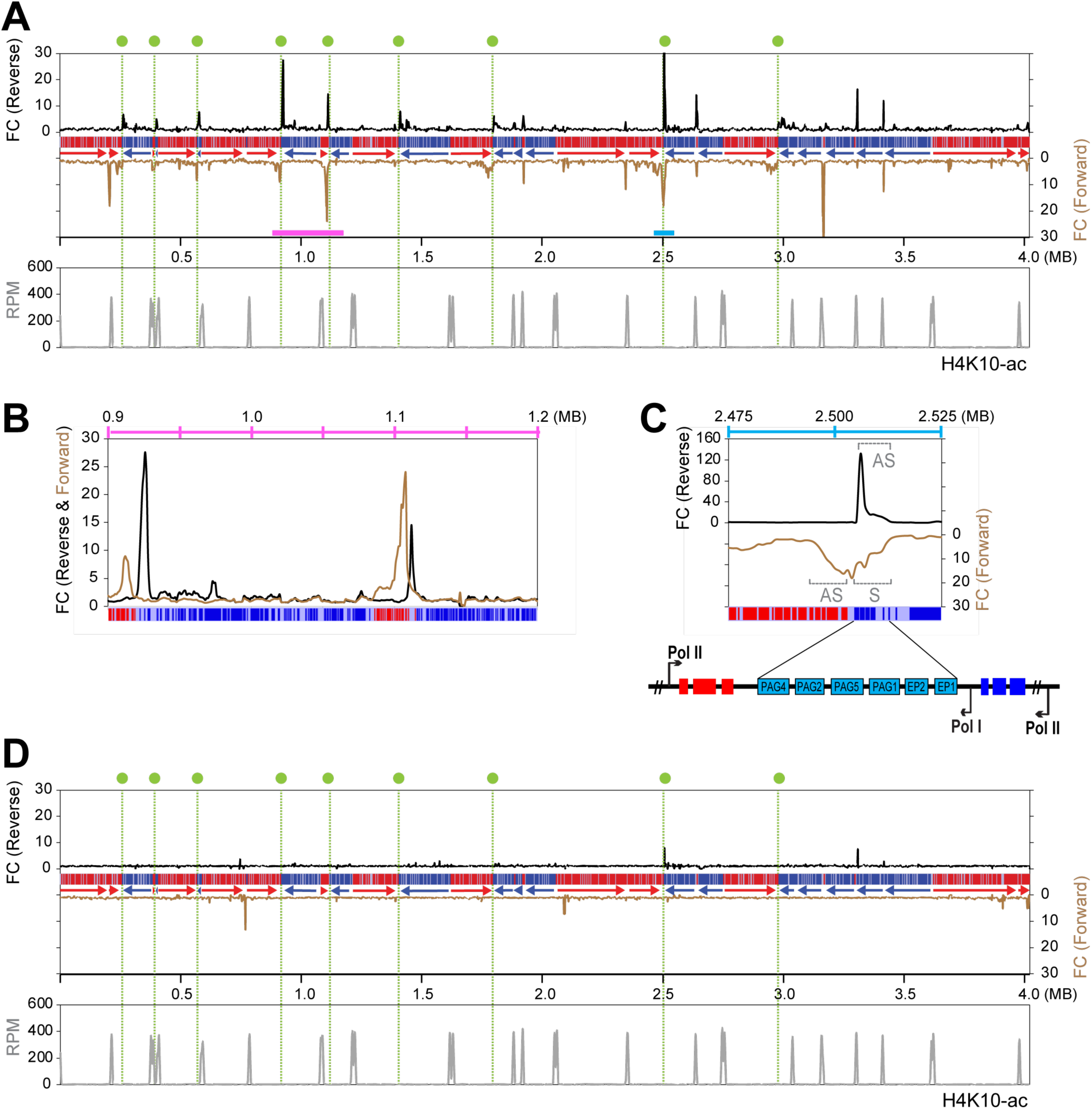
*Tb*FOP overexpression increases antisense transcription across TTSs. **(A)** Antisense transcription is increased near TTSs in *Tb*FOP-overexpressing cells. Reads from stranded RNA-seq were aligned to the *T. brucei* Lister 427 genome using Bowtie 2. Forward reads (secondary *y*-axis, brown line) and reverse reads (primary *y*-axis, black line) were analyzed separately using 5-kb sliding windows with a 1-kb step size. Fold changes in RPM values in *Tb*FOP OE cells relative to WT cells were plotted across all chromosomes. Chromosome 10 is shown as an example; the remaining chromosomes are shown in Fig. S5. Polycistronic transcription units (PTUs) are represented by red and blue bars, with arrowheads indicating the direction of transcription. Green dotted lines and circles indicate convergent transcription termination sites (TTSs). H4K10ac peaks marking transcription start sites (TSSs) are shown in the lower panel. Raw H4K10ac ChIP-seq reads from reference [11] were aligned to the Lister 427 genome using Bowtie 2. **(B)** Enlarged view of the 0.9–1.2-Mb region containing two convergent TTSs. **(C)** A Pol I-transcribed PTU containing procyclin genes and procyclin-associated genes (PAGs) in the 2.475–2.525-Mb region. The diagram illustrates the Pol I- and Pol II-transcribed PTUs and their transcriptional orientations. AS, antisense transcription; S, sense transcription. **(D)** *Tb*FOP depletion does not substantially alter genome-wide transcript patterns. Sequence reads were analyzed as described for panel A.

*Tb*FOP OE caused a pronounced increase in transcript levels throughout the genome at convergent TTSs and at some TTSs between head-to-tail PTUs (Fig. 5A). Two convergent TTSs within the 0.9–1.2-Mb region indicated by the pink bar in Fig. 5A are shown at higher resolution in Fig. 5B. The elevated signals reflected increased antisense transcription immediately upstream (brown line; consistent with readthrough from the reverse PTU) or downstream (black line; consistent with readthrough from the forward PTU) of the TTSs. Similar patterns of increases in antisense transcript levels were observed on other chromosomes (Fig. S5). Transcript levels at divergent transcription start sites (TSSs) remained largely unchanged following *Tb*FOP OE.

Given that *Tb*FOP OE increased the expression of Pol I-transcribed ES-associated VSGs (Fig. 4), we examined another Pol I-transcribed locus in the core region of chromosome 10 (Fig. 5C). This PTU, containing procyclin genes (EPs) and procyclin-associated genes (PAGs), is located in the 2.475–2.525-Mb region indicated by the blue bar in Fig. 5A. These PF stage-specific genes are normally repressed in BF trypanosomes [52]. Analysis of reverse reads (black line) revealed increased antisense transcripts across the procyclin PTU (marked AS). Analysis of forward reads (brown line) likewise revealed increased antisense transcription across the adjacent forward Pol II PTU (marked AS), consistent with transcriptional readthrough at the intervening TTS. Sense transcript levels across the procyclin Pol I PTU were also increased (brown line; marked S). This increase could reflect readthrough from the upstream Pol II PTU or altered regulation of life-cycle-specific procyclin transcription following *Tb*FOP OE. In contrast, *Tb*FOP depletion did not substantially alter genome-wide transcript patterns (Fig. 5D).

### *Tb*FOP overexpression preferentially affects promoter-proximal regions of silent BESs

Given that *Tb*FOP OE affected MES VSGs more strongly than BES VSGs (Fig. 4), we asked whether the effects of *Tb*FOP OE were regionally distributed within BESs. Reverse reads, representing sense transcripts, mapping to thirteen BESs were analyzed using 5-kb sliding windows with a 1-kb step, and RPM values from WT and *Tb*FOP OE cells were plotted across all BESs (Fig. 6A). *Tb*FOP OE strongly increased sense transcript levels in promoter-proximal regions of most silent BESs, whereas telomere-proximal regions were much less affected (Fig. 6A).

**FIG 6.**
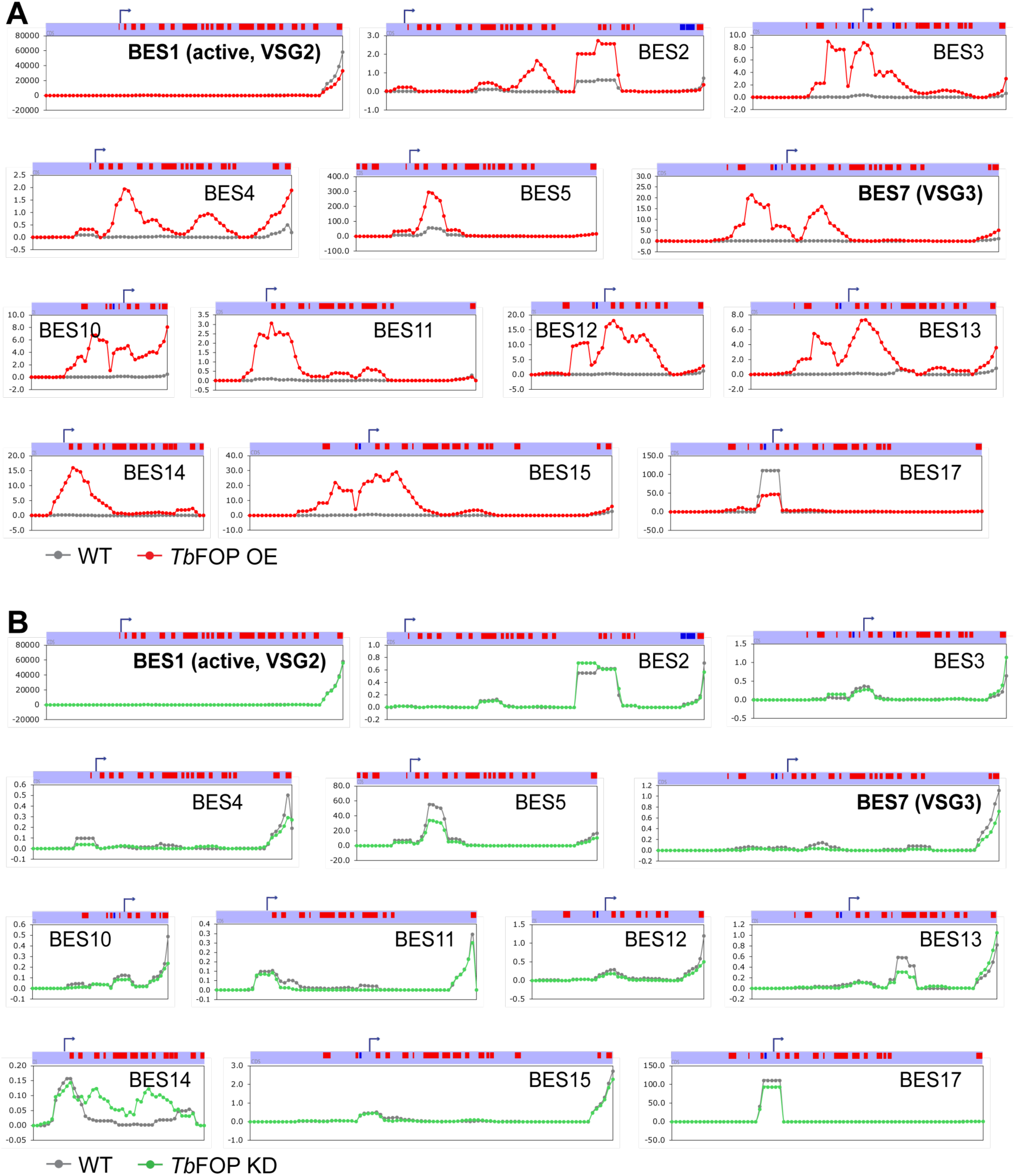
*Tb*FOP overexpression preferentially affects promoter-proximal regions of silent BESs. **(A)** Effect of *Tb*FOP overexpression on transcript levels across BESs. Forward and reverse reads mapping to 13 BESs were analyzed separately using 5-kb sliding windows with a 1-kb step. RPM values obtained from reverse reads, representing sense transcripts, were compared between WT (gray circles and lines) and *Tb*FOP OE (red circles and lines) cells. Forward reads, representing antisense transcripts, are presented in Fig. S6. Bent arrows indicate RNA Pol I promoters and the direction of transcription, and red boxes represent genes transcribed from left to right. **(B)** Effect of *Tb*FOP depletion on transcript levels across BESs. Sequence reads were analyzed as described for panel A. Green circles and lines represent values obtained from *Tb*FOP KD cells.

Two scenarios could account for this pronounced regional derepression. First, *Tb*FOP OE may perturb heterochromatin-mediated silencing near BES promoters, thereby allowing increased local transcription. Alternatively, Pol II transcription originating in subtelomeric or chromosome core regions may extend through transcription termination sites into silent BES PTUs. Consistent with the latter possibility, *Tb*FOP OE broadly increased transcripts associated with subtelomeric genes in both annotated orientations. Among 6,208 subtelomeric genes, 539 showed a greater than 2-fold increase (*P* < 0.01) in sense transcripts, and 369 showed a greater than 2-fold increase (*P* < 0.01) in antisense transcripts (Fig. S7 and Tables S9–S11). Because transcriptional organization and directionality within subtelomeric regions are poorly defined, the origins of these transcripts remain unclear. Nevertheless, their broad increase suggests extensive transcriptional derepression that could contribute to increased transcription in BES promoter-proximal regions. A similar pattern of transcriptional readthrough into BESs was observed in *T. brucei* cells lacking PP1-1 [23], suggesting a possible link between *Tb*FOP and the PJW/PP1 complex at these loci.

Analysis of forward reads further showed that *Tb*FOP OE also increased antisense transcript levels in BES promoter-proximal regions (Fig. S6A), raising the possibility that some transcripts originate downstream of BES promoters and proceed in the antisense direction. In contrast, *Tb*FOP depletion did not substantially affect BES silencing (Fig. 6B, Fig. S6B, and Tables S9–S11).

Because base J is strongly reduced at PTU boundaries and telomeric regions in PF trypanosomes [14, 15, 53], and *Tb*FOP OE alters transcription at J-enriched loci, we sought to explore the effects of *Tb*FOP OE by introducing the *Tb*FOP OE construct into PF cells. However, *Tb*FOP overexpression was not achieved following tetracycline induction using the same *Tb*FOP OE construct employed in BF cells (Fig. S8). Therefore, the potential dependence of the *Tb*FOP OE phenotype on base J could not be assessed in PF cells.

## DISCUSSION

*Tb*FOP overexpression produces a pronounced transcriptional phenotype consistent with defective transcription termination in *T. brucei*. The phenotypes observed in *Tb*FOP-overexpressing cells resemble those of mutants lacking TTS-associated chromatin marks (H3v, H4v, and base J), including cell lethality, abnormal increases in the G2/M and sub-G1 cell populations, and increased antisense transcription [19]. However, unlike these TTS mutants, which exhibit increased antisense transcription at both TTSs and TSSs, *Tb*FOP overexpression preferentially increased antisense transcription at TTSs. These similarities and differences suggest that *Tb*FOP may influence chromatin-associated processes at TTSs.

Another possibility is that the excess *Tb*FOP interferes with the PJW/PP1 complex or the base J biosynthesis machinery, thereby producing a similar transcriptional phenotype. Base J synthesis involves the generation of 5hmU by JBP1 and JBP2, followed by its glucosylation by JGT [14, 15]. JBP3, another J-binding protein identified through affinity purification of JGT in both *Leishmania* and *T. brucei*, is also required for transcription termination [20, 21]. Depletion of JBP3 results in transcriptional readthrough at termination sites and increased antisense transcription downstream of TTSs [20, 21]. The interaction between JBP3 and PNUTS is required for the recruitment of the PJW/PP1 complex to PTU boundaries [21–23, 54], and depletion of other components of this complex also causes transcription termination defects. Notably, depletion of these termination factors also leads to aberrant transcription at TSSs, although the effects are generally less pronounced than those at TTSs. Thus, the preferential effect of *Tb*FOP OE at TTSs suggests that the consequences of *Tb*FOP overexpression differ from, or may be more restricted than, those caused by direct disruption of known termination factors.

The transcriptional phenotype associated with *Tb*FOP overexpression may also relate to the previously proposed involvement of FOP proteins in mRNA export. *Tb*FOP is an arginine-rich protein containing a conserved FOP domain. Human FOP, also known as CHTOP, participates in mRNA export through the TREX complex, specifically by interacting with UAP56, an RNA helicase component of TREX. Similarly, *T. cruzi* FOP has been shown to interact with *Tc*SUB2 (UAP56 ortholog) of the TREX complex [40, 55]. Although the overall sequence similarity between *Tb*FOP and human FOP/CHTOP is limited, these observations raise the possibility that *Tb*FOP may participate in RNA-associated processes that are functionally coupled to transcription. However, because depletion of *Tb*FOP did not produce detectable effects on global mRNA export in a previous study [40], its physiological role in these processes remains unresolved.

Human FOP has been reported to bind 5-hydroxymethyl cytosine (5hmC), an interaction important for its transcriptional regulatory activity [56]. This finding raises the possibility that *Tb*FOP could interact directly or indirectly with 5hmU/base J-associated chromatin at TTSs. Such an interaction could provide a link between *Tb*FOP abundance and transcription termination at base J-enriched TTSs. However, direct binding of *Tb*FOP to 5hmU or base J has not been demonstrated, and whether *Tb*FOP influences the localization or activity of JGT, JBP3, or other components of the PJW/PP1 complex remains to be determined. Our attempt to test whether the *Tb*FOP OE phenotype depends on base J by expressing *Tb*FOP in PF cells was inconclusive because efficient *Tb*FOP overexpression could not be achieved in PF cells.

*Tb*FOP overexpression also altered BES expression in a strongly position-dependent manner, with the largest increases in transcript levels occurring near silent BES promoters. One possible explanation is that transcription from the terminal Pol II-transcribed PTU in the chromosome core may read through its termination site and extend toward the silent BES, similar to the transcriptional readthrough observed in *T. brucei* cells lacking PP1-1 [23]. Alternatively, transcription may initiate nonspecifically within subtelomeric regions and extend into downstream BESs. The levels of many subtelomeric transcripts increased in both the sense and antisense directions in *Tb*FOP OE cells, suggesting extensive transcriptional derepression within subtelomeric regions that are normally repressed.

Several observations indicate that the phenotypes described here should be interpreted specifically as consequences of *Tb*FOP overexpression rather than as evidence that endogenous *Tb*FOP is required for transcription termination or VSG silencing. *Tb*FOP depletion did not substantially affect cell growth, global transcription patterns, or VSG expression under the conditions examined. Moreover, *Tb*FOP overexpression causes severe cytotoxicity, and some transcriptional or cell-cycle changes may therefore represent secondary consequences of cellular dysfunction. Nevertheless, the preferential accumulation of antisense transcripts at TTSs, rather than a comparable increase at TSSs or throughout PTUs, suggests that the transcriptional phenotype is not simply a nonspecific consequence of toxicity.

Together, our results show that increased *Tb*FOP levels strongly perturb transcription at TTSs and disrupt VSG silencing across multiple genomic contexts. Determining whether *Tb*FOP directly interacts with TTS-associated chromatin or transcription termination factors, and establishing its physiological function under endogenous expression conditions, will be important for understanding the mechanistic basis of these phenotypes. Such studies may provide new insights into the unusual mechanisms that regulate transcription termination and VSG silencing in *T. brucei*.

## MATERIALS AND METHODS

### Trypanosoma brucei cell lines

Bloodstream form *Trypanosoma brucei* cells were cultured in HMI-9 medium supplemented with 10% fetal bovine serum (FBS; Cytiva HyClone) at 37°C with 5% CO_2,_ as described previously [57]. Necessary antibiotics (InvivoGen) were added at the following concentrations: G418 at 2.5 µg/ml, puromycin at 0.1 µg/ml, phleomycin at 1 µg/ml, blasticidin at 5 µg/ml, and hygromycin at 5 µg/ml. Cell lines used in this study are listed in Table S1. All strains were generated in the AMT2 WT strain (the 2T1 background) [51, 58]. The strain carries the tetracycline-inducible system with T7 RNA polymerase and a Tet repressor (TetR) to control the expression of a gene of interest or of double-stranded RNA (dsRNA) for target mRNA depletion.

Procyclic form *Trypanosoma brucei* cells in the PF29-13 background [59] carrying the Tet-inducible system were cultured in SDM-79 medium supplemented with 10% FBS (Cytiva HyClone) and 7.5 µg/ml hemin (Sigma) at 28°C. Necessary antibiotics (InvivoGen) were added at the following concentrations: G418 at 15 µg/ml, hygromycin at 50 µg/ml, and blasticidin at 20 µg/ml.

*Trypanosoma brucei* cell lines, plasmids, and oligonucleotides (Integrated DNA Technologies, IDT) used in this study are listed in Tables S1, S2, and S3, respectively.

### Flow cytometry

Five million cells were fixed in ice-cold 75% ethanol and incubated with 50 µg/ml propidium iodide (Sigma) and 200 µg/ml RNase A (Sigma) in 1x PBS (Corning) at 37°C for 30 min in the dark. Samples were analyzed for cell-cycle progression using flow cytometry (FACSVia, BD Biosciences), FlowJo, and FlowJo models Watson Pragmatic.

To detect GFP-expressing cells, live BF trypanosomes collected on days 0, 1, and 2 after tetracycline addition were analyzed by flow cytometry.

### Immunoblotting

Cells were harvested at the indicated time points and suspended in 2x Laemmli buffer (Bio-Rad) at a density of 0.25 million cells per µl. Denatured proteins from whole cells were separated on a 4-20% gradient SDS-PAGE gel (Bio-Rad) and transferred to a nitrocellulose membrane (GE Healthcare). The membranes were probed in 1x PBS supplemented with 0.1% Tween 20 and 1% milk (Bio-Rad) with the following antibodies: anti-6xHis (monoclonal mouse, Sigma, H1029, 1:1,000 dilution), HA (polyclonal rabbit, Sigma, H6908, 1:2,000 dilution), GFP (polyclonal rabbit, Sigma, G1544, 1:1,000 dilution), histone H3 (monoclonal rabbit, ABclonal, AP84276, 1:2,000 dilution), or VSG3 (polyclonal rabbit, a gift from George Cross’s lab, 1:5,000 dilution). The membranes incubated with a primary antibody overnight at 4°C were washed with 1x PBS supplemented with 0.1% Tween 20 and then incubated with secondary antibodies: rabbit anti-IgG conjugated with Horseradish Peroxidase (HRP) (GE Healthcare, NA934V, 1:5,000 dilution) or mouse anti-IgG conjugated with HRP (GE Healthcare, NA931V, 1:5,000 dilution). The membranes were sprayed with WesternBright ECL solutions (Thomas Scientific, C942A86), and corresponding proteins were detected using the ChemiDoc Imaging System (Bio-Rad).

### Overexpression library screen

The pRPa-Sce* vector contains a Tet-inducible endonuclease I-SceI gene and its 18-bp recognition sequence [32]. In the AMT2 strain, the landing pad locus, an rDNA spacer region marked by the puromycin-resistance gene (PUR), was targeted with a linearized pRPa-Sce* vector. This targeting replaced the PUR marker with the I-SceI gene and its 18-bp recognition site, generating the HSTB-1297 strain. The HSTB-1297 cells were treated with 1 µg/ml tetracycline for 3 hours to introduce a double-strand break at the landing pad and then transfected with about 100 µg of linearized *T. brucei* ORF library vectors to integrate individual ORFs into the DSB site. Approximately 400,000 independently targeted ORF clones were obtained.

To induce ORF overexpression, the *T. brucei* ORF libraries were treated with or without 2 µg/ml tetracycline for three days in triplicate. Genomic DNA was isolated from each culture (three uninduced and three Tet-induced) using the QIAamp Blood Mini Kit (Qiagen, 51104) and sonicated with Bioruptor Pico (Diagenode) with 5–7 cycles of 30 seconds on/off. DNA (1.5 µg) was then processed for next-generation sequencing (NGS) library preparation using the NEBNext Ultra II DNA Library Prep Kit (NEB, E7645S), NEBNext Multiplex Oligos (NEB, E7335S), and AMPure XP beads (Beckman, A63881). Blunted and adaptor-ligated DNA was amplified using a custom forward oligo (HK898), which anneals to the attB1 site of the ectopically integrated ORF, and an indexed universal reverse primer. The NGS libraries were sequenced on an Illumina HiSeq 4000 using a custom oligo (HK899) annealing to the attB1 site, with a 150-bp single-end (SE) configuration.

Read quality was analyzed using FastQC, and reads were trimmed using Trim Galore (parameters: phred 33, stringency 1 bp, error 0.1, length 20) and aligned to the *Tb*927v5 genome using Bowtie 2 with default parameters. This reference genome was used for mapping because it was used to design the oligos for the ORF library [31]. Aligned reads were analyzed using SeqMonk (MAPQ >20 filter) and DESeq from Babraham Bioinformatics. The alignment report, correlations between replicates, and the PCA analyses are summarized in Table S4.

### Stranded RNA-seq with rRNA depletion

WT (AMT2) and *Tb*FOP OE (HSTB-1461) strains were treated with 1 µg/ml tetracycline for one day, and the *Tb*FOP KD (NPT-95) strain was treated for two days. Cultures were prepared in triplicate. Total RNA was extracted from approximately 50 million cells using RNA Stat-60 (Tel-Test) according to the manufacturer’s protocol, cleaned with an RNeasy kit (Qiagen), and quantified with a NanoDrop 2000c spectrophotometer (Thermo Fisher Scientific). Ribosomal RNA was depleted by Novogene using a proprietary method. Stranded RNA-seq libraries were prepared using random hexamers and the NEBNext Ultra II Directional RNA Library Prep Kit for Illumina (NEB, E7760) and sequenced on a NovaSeq X Plus.

Read quality was analyzed using the FastQC program, and reads were trimmed using the Trim Galore program (parameters: phred 33, stringency 1 bp, error 0.1, length 20) and aligned to the Lister 427 genome (v66) and VSGnome using Bowtie 2 [6, 29]. Aligned reads were analyzed using SeqMonk (MAPQ >20 filter). The alignment report, correlations between replicates, and the PCA analyses are summarized in Tables S6 and S7.

## DATA AVAILABILITY

The raw NGS data from the OE library screen and stranded RNA-seq have been deposited in the NCBI Sequence Read Archive under BioProject accession number PRJNA1313143.

## SUPPLEMENTARY DATA

Supplementary data are available online.

## AUTHOR CONTRIBUTIONS

H.K. and E.E. conceived and designed the study. N.P.S. and A.M. generated plasmids and cell lines, performed candidate validation experiments for BF *T. brucei*, and participated in the OE screen and RNA-seq experiments. V.G. performed the growth and immunoblotting experiments for PF *T. brucei*. E.E. provided the vectors used to construct the *T. brucei* ORF library and independently analyzed the OE screen NGS data. H.K. performed the OE screen and targeted NGS, analyzed the OE screen NGS and RNA-seq data, prepared the figures, and supervised the study. H.K. wrote the original draft. All authors reviewed and revised the manuscript and approved the final manuscript.

## ACKNOWLEDGEMENTS

We thank Azenta for Illumina sequencing of the OE library screen samples, Novogene for stranded RNA-seq library preparation and Illumina sequencing, and the PHRI Flow Cytometry Core for its services. ChatGPT (OpenAI) was used solely for language editing and to improve the clarity and readability of author-generated text. All suggestions were reviewed and revised by the authors.

