## Supplementary Information for "A genome-wide overexpression screen identifies *Tb*FOP, whose overexpression disrupts transcription termination in *Trypanosoma brucei*"

### Supplemental Tables

**Table S1.** ***Trypanosoma brucei*** **cell lines used in this study**

| Name | Genotype | Reference |
|---|---|---|
| 2T1 | PUR at rDNA locus (landing pad) in 2T1 strain (T7 RNA Polymerase & Tet Repressor with PHLEO marker) | [1] |
| AMT2 | PHLEO marker replaced with NEO in the 2T1 strain | [2] |
| HSTB-1297 | AMT2 transfected with pRPa-Sce* | This study |
| HSTB-1298 | HSTB-1297 strain transfected with ORF library vectors | This study |
| HSTB-1283 | AMT2 strain with a silent BES promoter marked with GFP-PHLEO | This study |
| HSTB-1461 | *Tb*FOP OE strain: Tet-inducible N-6xHis-*Tb*FOP cassette (BSD marker) integrated at the landing pad locus in HSTB-1283 | This study |
| HSTB-1466 | *Tb*927.5.1270 OE strain: Tet-inducible N-6xHis-*Tb*927.5.1270 cassette (BSD marker) integrated at the landing pad locus in HSTB-1283 | This study |
| HSTB-1469 | *Tb*927.11.2250 OE strain: Tet-inducible N-6xHis-*Tb*927.11.2250 cassette (BSD marker) integrated the landing pad locus in HSTB-1283 | This study |
| NPT81 | *Tb*FOP-3xHA-BSD/WT: One *Tb*FOP allele is tagged with 3xHA at the C-terminus in AMT2 strain | This study |
| NPT95 | *Tb*FOP RNAi KD: *Tb*FOP-HA-BSD/WT (NPT81) transfected with *Tb*FOP RNAi KD vector (HYG marker) | This study |
| PF29-13 | Procyclic form WT strain with T7 RNA Polymerase & Tet Repressor with NEO and HYG markers | [3] |
| VTB101 | Procyclic form *Tb*FOP OE strain: Tet-inducible N-6xHis-*Tb*FOP cassette (BSD marker) integrated at the rDNA locus in PF29-13. | This study |

**Table S2. Plasmids used in this study**

| Name | Description | Reference |
|---|---|---|
| p2T7-NEO | T7 RNA polymerase, NEO marker | [2] |
| p2T7-TA-HYG | RNAi KD vector | [4] |
| pSUN48 | GFP-PHLEO with BES promoter targeting sequences | This study |
| pSUN87 | Tet-inducible vector with an N-terminal 6xHis-tag and the rDNA landing pad targeting sequence (BSD marker) | This study |
| pOE5 | N-6xHis-*Tb*927.5.1270 ectopic expression vector with the rDNA landing pad targeting sequence (in pSUN87, BSD) | This study |
| pOE6 | N-6xHis-*Tb*FOP ectopic expression vector with the rDNA landing pad targeting sequence (in pSUN87, BSD) | This study |
| pOE8 | N-6xHis-*Tb*927.11.2250 ectopic expression vector with the rDNA landing pad targeting sequence (in pSUN87, BSD) | This study |
| pNPS27 | *Tb*FOP RNAi KD vector (p2T7-TA-HYG) | This study |
| pMOTag-6H | C-terminal in-situ tagging vector with 3xHA and BSD marker | This study |

**Table S3. Oligonucleotides used in this study**

| Name | Description | Sequences |
|---|---|---|
| NP 9 | Forward to amplify *Tb*927.5.1270 CDS | ATGGAGCTTCTCAAGAAAACGG |
| N10 | Reverse to amplify *Tb*927.5.1270 CDS | TTCCGTTCGCACAACCTTAGA |
| NP11 | Forward to amplify *Tb*927.6.1470 (*Tb*FOP) CDS | ATGCGCCGAGACGACTTC |
| NP12 | Reverse to amplify *Tb*927.6.1470 (*Tb*FOP) CDS | AGCCGCCCCTCGGAACCG |
| NP15 | Forward to amplify *Tb*927.11.2250 CDS | ATGTACAATAACGACTCGGCG |
| NP16 | Reverse to amplify *Tb*927.11.2250 CDS | GGAAGGCCCATCTGCCTTG |
| HK1333 | Forward to tag *Tb*927.6.1470 (*Tb*FOP) with 3xHA using pMOTag-6H | aaaagaaggggctcgcagaagtgggcccatgacgcgc gagggtcttgatgagcagttggatcggttccgaggggcgg ct GGTACCGGGCCCCCCCTCGAG |
| HK1334 | Reverse to tag *Tb*927.6.1470 (*Tb*FOP) with 3xHA using pMOTag-6H | actcttttgttatggtggtagaagtcccatttccgcccttcaa cagccttcagagtttcaaaactactcctcca TGGCGGCCGCTCTAGAACTAGTGGAT |
| HK-898 | Custom forward primer to amplify ectopic ORFs in NGS library preparation | aatgatacggcgaccaccgagatctacaccctgcagggg gacaagtttgtacaaaaaagcaggct |
| HK-899 | NGS sequencing primer for ORF identification from the OE screen | gggggacaagtttgtacaaaaaagcaggctATG |

**Table S4. OE toxicity screen NGS alignment summary (PCA, replicate correlations, and coverage)**

**Table S5. Toxic gene identification by targeted NGS.** (A) Raw counts from reads aligned to the *Tb*927v5 genome. (B) DESeq-analyzed genes. (C) DESeq-analyzed genes that are included in the original ORFeome library. (D) Underrepresented ORFs with *P* adj < 0.1. (E) Over-represented ORFs with *P* adj < 0.1. (F) ORFs that were not recovered from this screen.

**Table S6. Stranded RNA-seq: VSGnome analysis summary.** (A) Log2(RPKM) values for VSG genes: WT vs OE in triplicate. (B) Log2(RPKM) values for VSG genes: WT vs KD in triplicate. (C) Summary of statistical analysis for boxplots using GraphPad Prism. (D) Alignment summary (PCA, replicate correlations, and coverage).

**Table S7. Stranded RNA-seq: Lister 427 genome alignment summary (PCA, replicate correlations, and coverage)**

**Table S8. Sliding window analysis of reads aligned to the Lister 427 genome.** Sequence reads from WT, *Tb*FOP OE, and *Tb*FOP KD samples in triplicate were analyzed. H4K10ac ChIP-seq reads from reference [5] were used to mark TSSs. (A) RPM values obtained from reverse reads aligned to the Lister 427 genome. (B) RPM values obtained from forward reads aligned to the Lister 427 genome.

**Table S9. Transcriptome analysis of all reads comparing WT, *Tb*FOP OE, and *Tb*FOP KD.** (A) All genes, OE vs WT. (B) Core genes, OE vs WT. (C) Subtelomeric genes, OE vs WT. (D) BES genes, OE vs WT. (E) All genes, KD vs WT. (F) Core genes, KD vs WT. (G) Subtelomeric genes, KD vs WT. (H) BES genes, KD vs WT.

**Table S10. Transcriptome analysis of sense reads comparing WT, *Tb*FOP OE, and *Tb*FOP KD.** (A) All genes, OE vs WT. (B) Core genes, OE vs WT. (C) Subtelomeric genes, OE vs WT. (D) BES genes, OE vs WT. (E) All genes, KD vs WT. (F) Core genes, KD vs WT. (G) Subtelomeric genes, KD vs WT. (H) BES genes, KD vs WT.

**Table S11. Transcriptome analysis of antisense reads comparing WT, *Tb*FOP OE, and *Tb*FOP KD.** (A) All genes, OE vs WT. (B) Core genes, OE vs WT. (C) Subtelomeric genes, OE vs WT. (D) BES genes, OE vs WT. (E) All genes, KD vs WT. (F) Core genes, KD vs WT. (G) Subtelomeric genes, KD vs WT. (H) BES genes, KD vs WT.

### Supplemental Figures

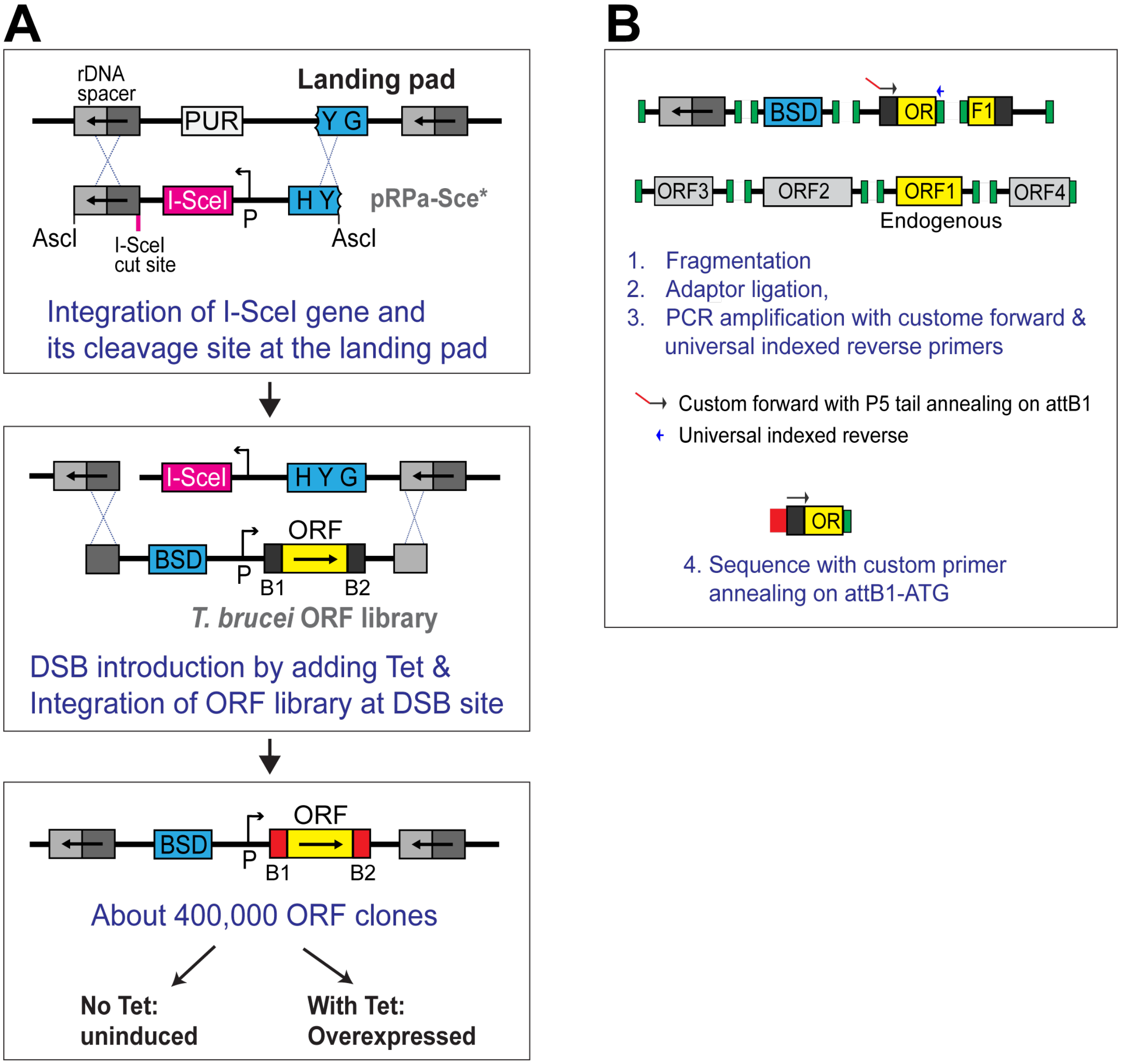


**FIG S1. OE-induced toxicity screening. (A)** Integration of the ORF library at the rDNA spacer landing pad (LP) locus. The AMT2 strain in the 2T1 background was transfected with AscI-digested pRPa-Sce* vector, resulting in integration of the Tet-inducible I-SceI gene, its cleavage site, and the HYG selection marker at the LP locus, replacing PUR and the truncated HYG gene. Transient Tet induction for 3 h generated a DSB at the LP locus, facilitating integration of the *T. brucei* ORF library. Library integration removed the DSB-inducible cassette. Approximately 400,000 ORF clones were obtained and screened for OE-induced toxicity by culturing cells with or without tetracycline for 3 days in three independent cultures. **(B)** Assessment of individual ORF representation by targeted next-generation sequencing (NGS). Genomic DNA was fragmented and ligated to an Illumina adaptor. ORFs were amplified using a custom p5 forward oligo annealing to the attB1 site and an indexed universal reverse oligo. The resulting NGS libraries were sequenced using a custom primer annealing at the attB1-ATG site.

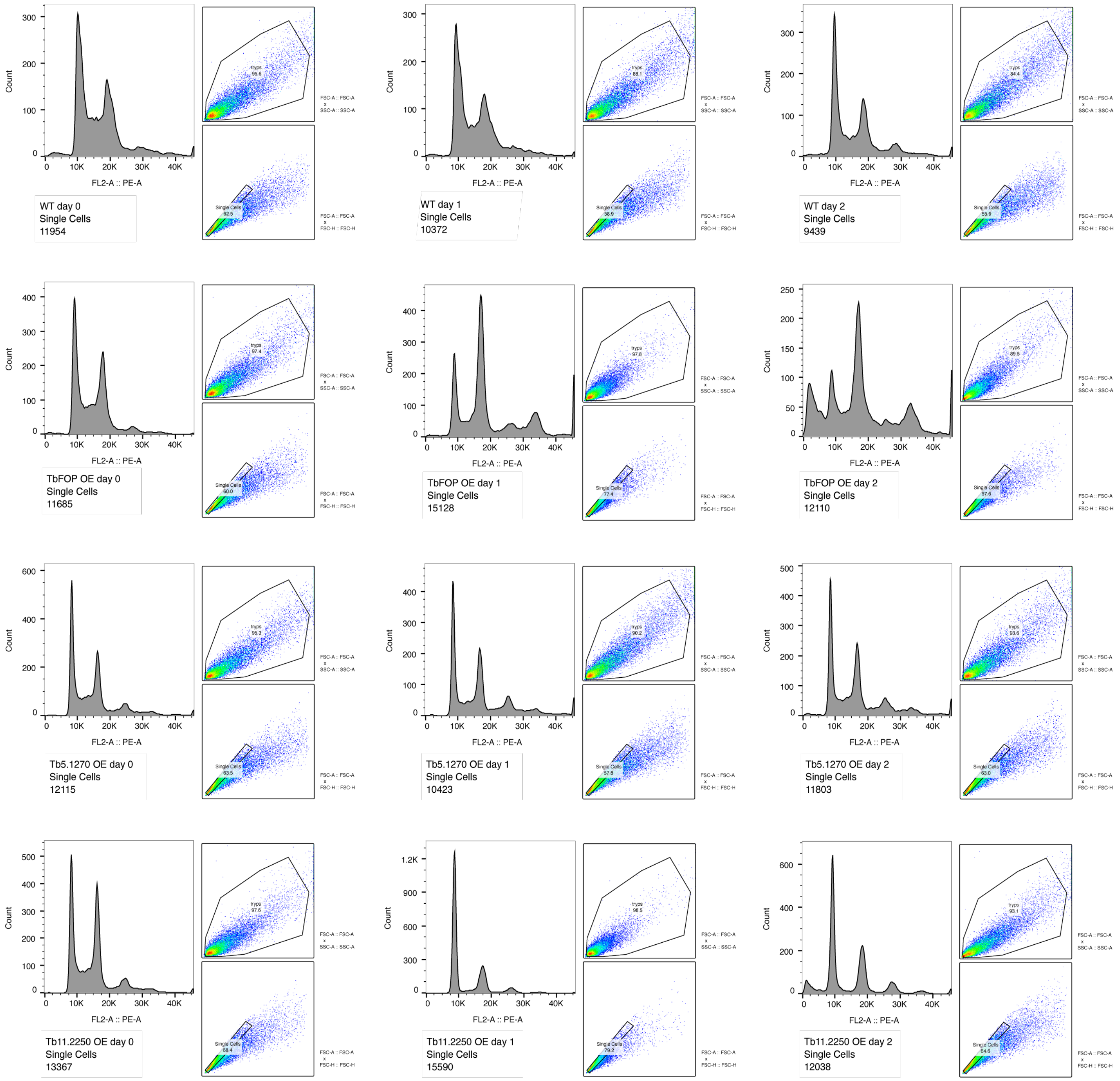


**FIG S2. Gating strategies used to analyze the cell-cycle profiles of WT or cells overexpressing *Tb*FOP, *Tb*927.5.1270, or *Tb*927.11.2250.** The two scatter plots adjacent to each cell-cycle histogram show gating of the trypanosome cell population (top) and single-cell population (bottom).

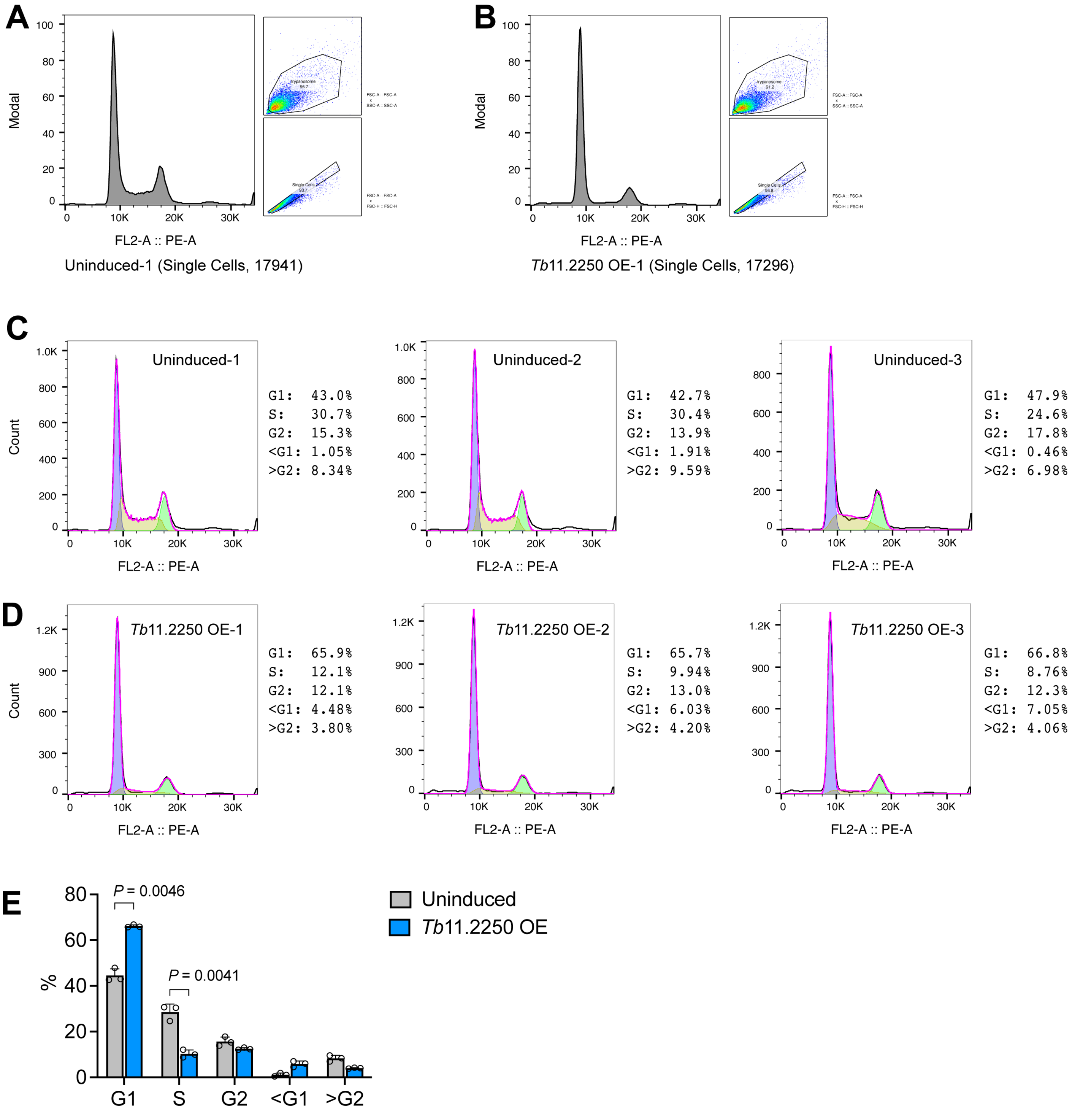


**FIG S3. *Tb*927.11.2250 overexpression causes a significant reduction in the S-phase population. (A, B)** Gating strategies shown for one replicate of WT and *Tb*927.11.2250-overexpressing cells. WT and OE strains treated with or without Tet in triplicate were fixed in ice-cold 75% ethanol, stained with PI, and analyzed by flow cytometry. The two scatter plots adjacent to each cell-cycle histogram show gating of the trypanosome cell population (top) and single-cell population (bottom). **(C)** Cell-cycle profiles of uninduced cells. **(D)** Cell-cycle profiles of *Tb*927.11.2250-overexpressing cells. Cell-cycle distributions were analyzed using the Watson Pragmatic model in FlowJo. **(E)** Percentages of cells in different cell-cycle phases. Statistical significance was determined using unpaired *t* tests with Welch's correction in GraphPad Prism.

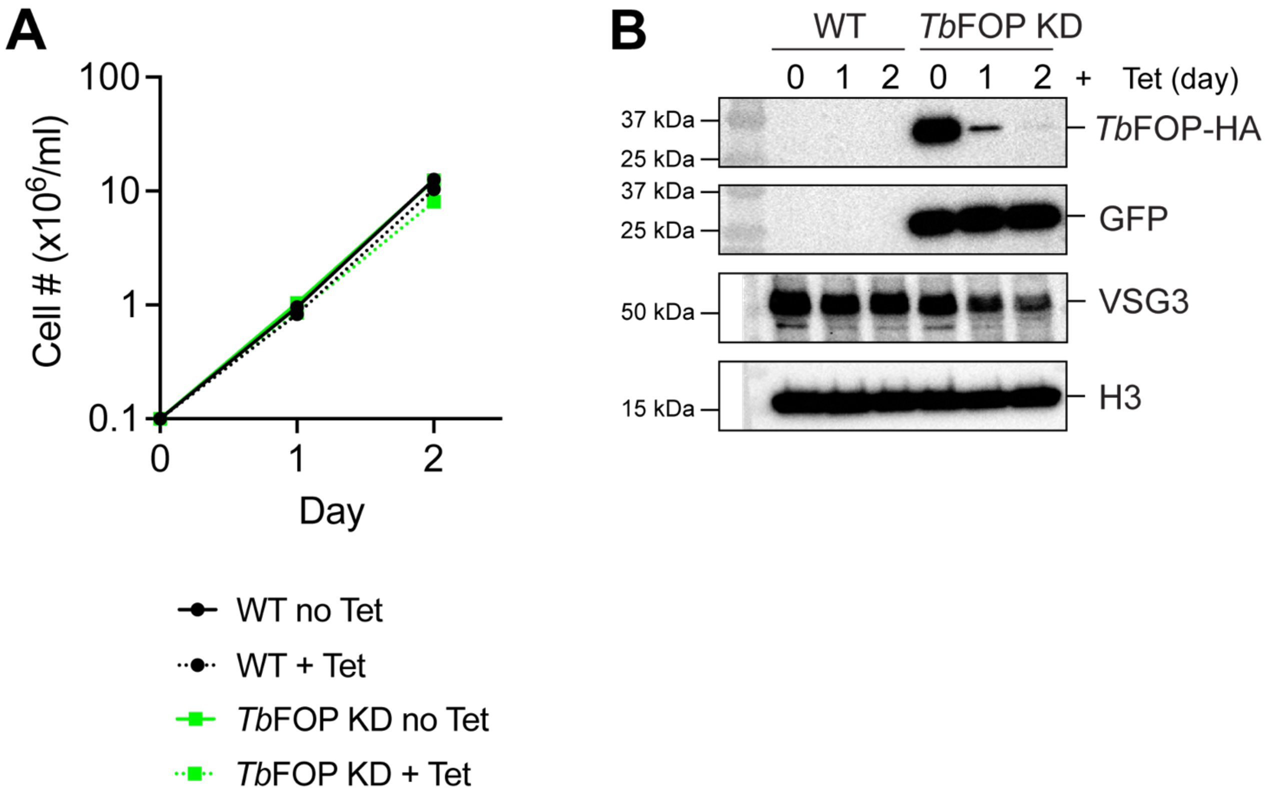


**FIG S4. *Tb*FOP depletion does not inhibit trypanosome cell growth. (A)** Effect of *Tb*FOP depletion on cell growth. The C-terminus of one *Tb*FOP allele in the WT *T. brucei* strain (AMT2) was epitope-tagged with 3xHA using pMOTag-6H (3xHA, BSD marker). The *Tb*FOP-HA strain was transfected with the Tet-inducible *Tb*FOP RNAi KD vector. Tetracycline was added to WT and *Tb*FOP KD cells, and cell growth was monitored daily for 2 days by counting cells. **(B)** Immunoblot confirming the depletion of *Tb*FOP protein. Proteins in whole cell extracts were analyzed by immunoblotting using antibodies against HA, GFP, VSG3, or histone H3 (loading control) antibodies.

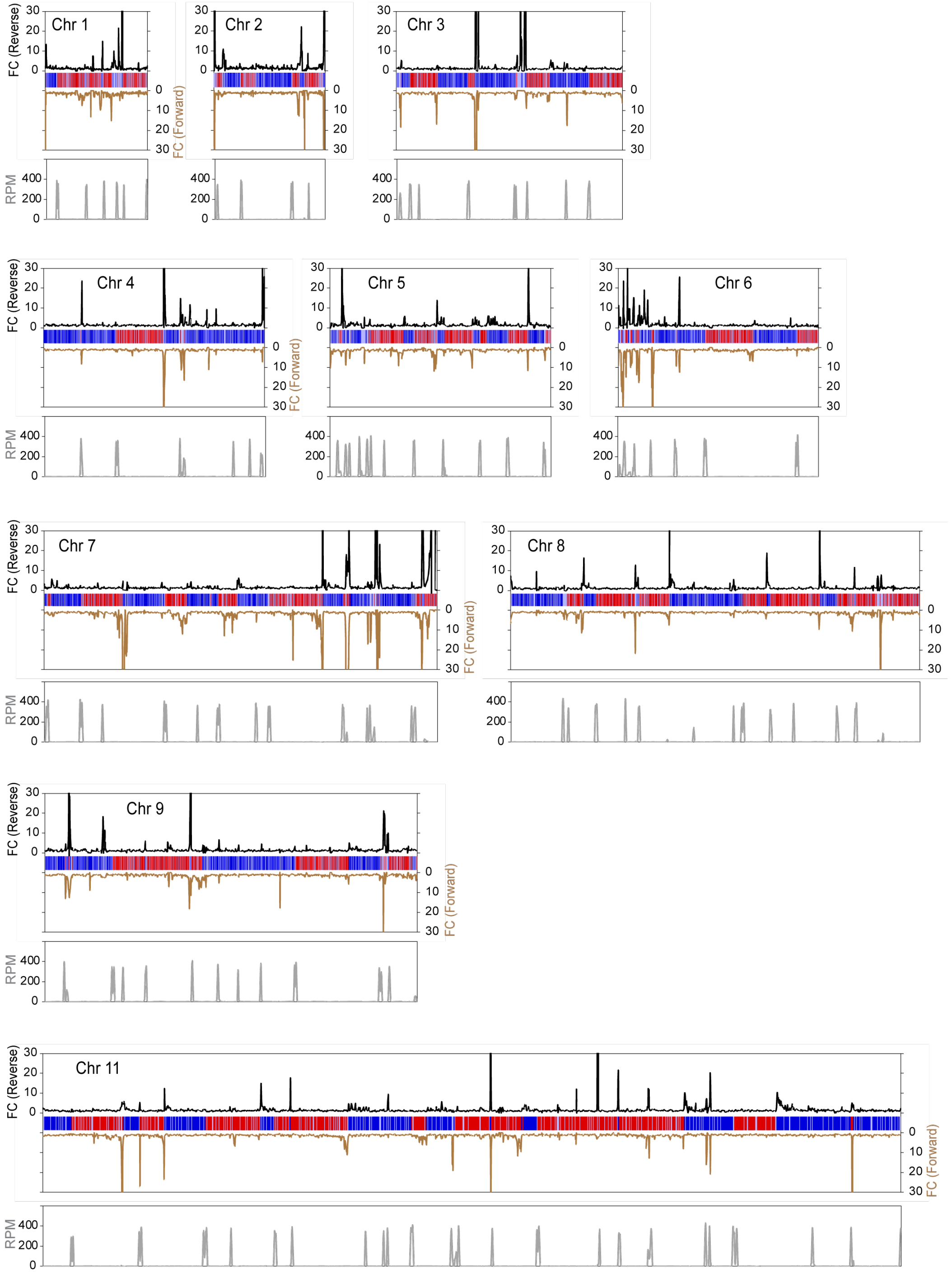


**FIG S5. *Tb*FOP overexpression increases antisense transcription across TTSs.** Reads from stranded RNA-seq were aligned to the *T. brucei* Lister 427 genome using Bowtie 2. Forward reads (secondary *y*-axis, brown line) and reverse reads (primary *y*-axis, black line) were analyzed separately using 5-kb sliding windows with a 1-kb step size. Fold changes in RPM values in *Tb*FOP OE cells relative to WT cells were plotted across all chromosomes. All chromosomes except chromosome 10 are shown. Polycistronic transcription units (PTUs) are represented by red and blue bars, with arrowheads indicating the direction of transcription. Green dotted lines and circles indicate convergent transcription termination sites (TTSs). H4K10ac peaks marking transcription start sites (TSSs) are shown in the lower panel. Raw H4K10ac ChIP-seq reads from reference [5] were aligned to the Lister 427 genome using Bowtie 2.

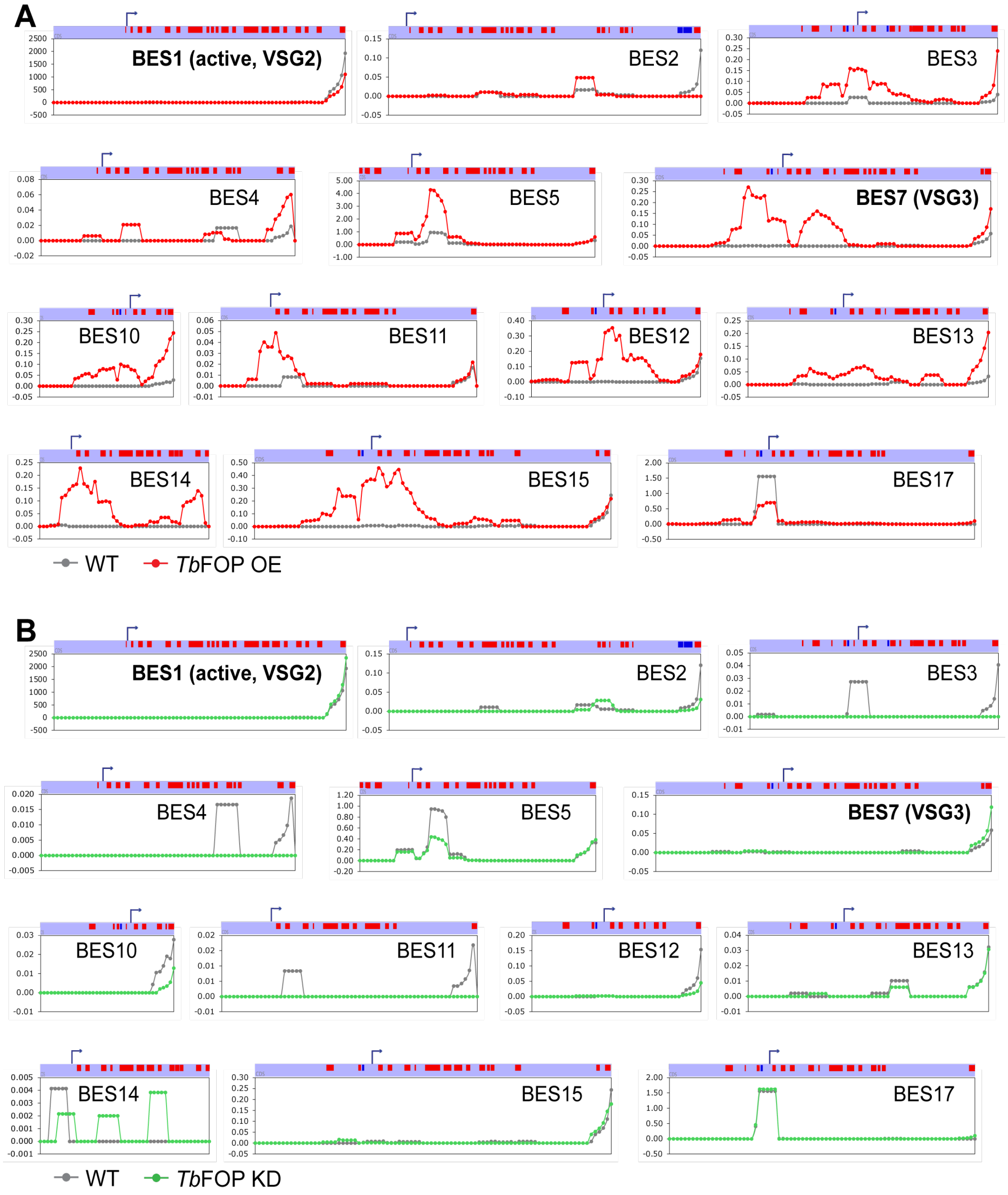


**FIG S6. Levels of antisense transcripts in BESs in TbFOP OE or KD cells.** Forward reads representing antisense transcripts were analyzed using 5-kb sliding windows with a 1-kb step. RPM values were compared **(A)** between WT (gray circles and lines) and *Tb*FOP OE (red circles and lines) cells or **(B)** between WT and *Tb*FOP KD cells (green circles and lines). Bent arrows indicate RNA Pol I promoters and the direction of transcription, and red boxes represent genes transcribed from left to right.

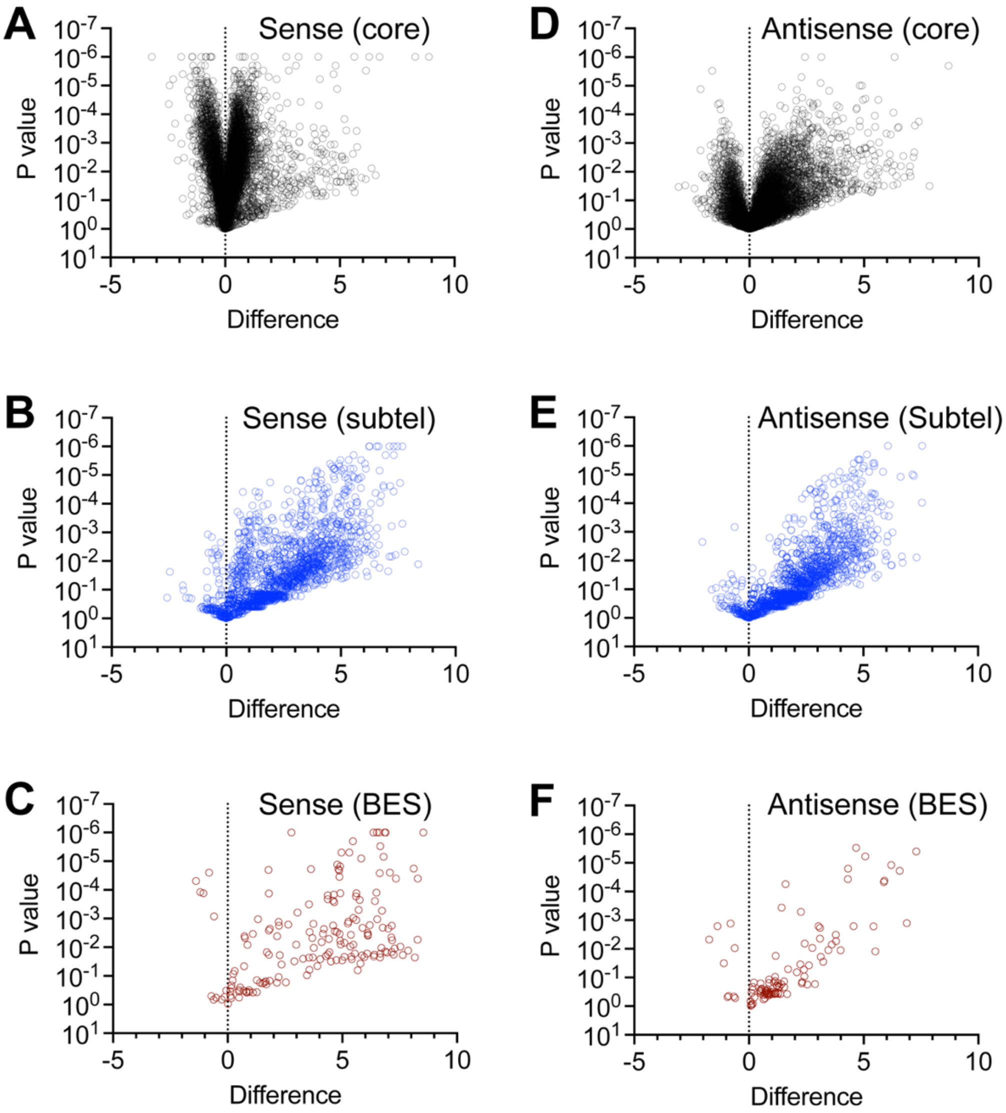


**FIG S7. Transcriptomic profiles of *Tb*FOP OE cells.** Sequence reads mapping to individual genes in the Lister 427 genome were analyzed separately for sense and antisense transcription. The log2(RPKM) values for individual genes were compared in scatter plots. Statistical significance was determined using multiple unpaired *t* tests with Welch's correction and the two-stage linear step-up procedure in GraphPad Prism. **(A)** Sense transcription of chromosome core genes. **(B)** Sense transcription of subtelomeric genes. **(C)** Sense transcription of BES genes. **(D)** Antisense transcription of chromosome core genes. **(E)** Antisense transcription of subtelomeric genes. **(F)** Antisense transcription of BES genes.

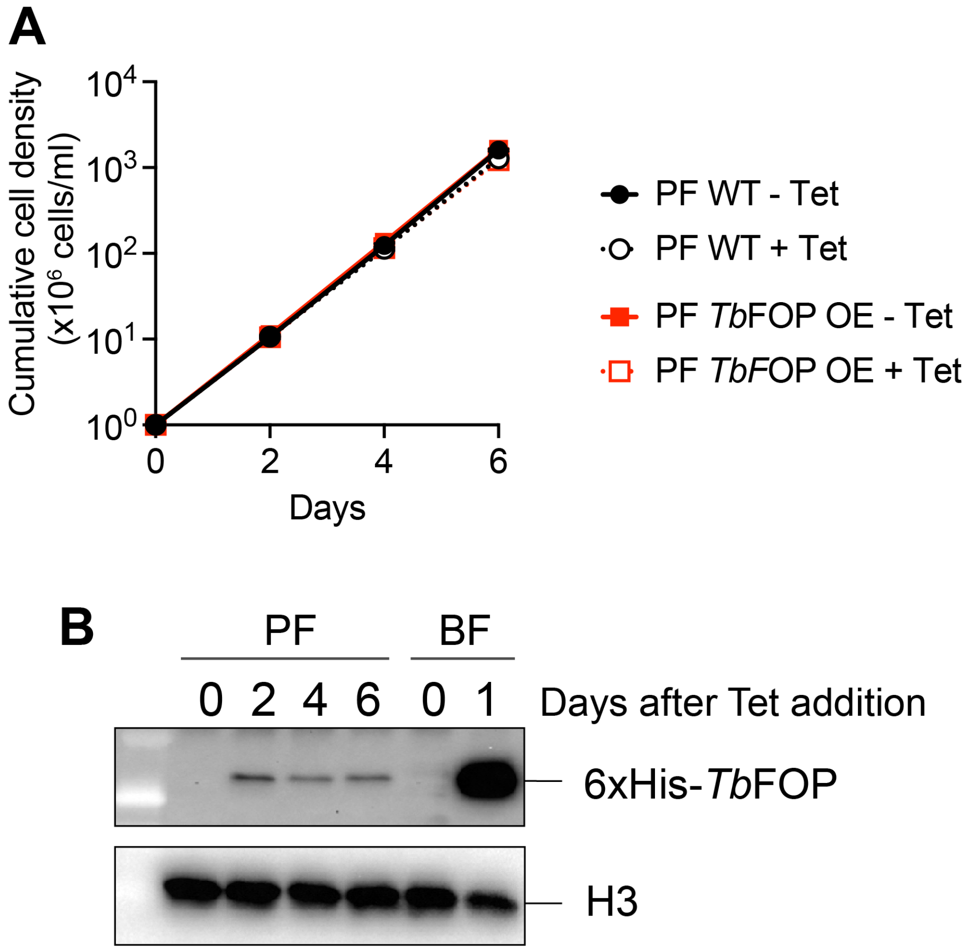


**FIG S8. *Tb*FOP overexpression is not achieved in procyclic-form cells. (A)** Wild-type and *Tb*FOP OE-inducible procyclic-form (PF) cells were cultured with or without tetracycline (Tet) for 6 days. Cell growth was monitored every 2 days by counting cells. **(B)** Expression of 6xHis-*Tb*FOP in PF cells at the indicated times after Tet addition. 6xHis-*Tb*FOP expression in bloodstream-form (BF) cells after 1 day of Tet induction is shown for comparison. Whole-cell lysates were analyzed by immunoblotting using antibodies against 6xHis and histone H3. Lysates corresponding to $2.5 \times 10^6$ cells were loaded per lane. Histone H3 served as a loading control.
